# Gene-level evolutionary conservation and human constraint decline at comparable rates across the age of onset of Mendelian disease

**DOI:** 10.64898/2026.10.08.757734

**Authors:** Anees Ahmed Mahaboob Ali, Everette Jacob Remington Nelson

## Abstract

**Background:** Gene-level constraint and conservation scores are widely used to prioritise disease genes and to benchmark genomic language models. Purifying selection acts weakly on phenotypes manifesting after reproduction, so these scores are expected to carry less information about late-onset disease. An onset-dependent gradient has been demonstrated for human allele frequencies but not for the gene-level scores used in practice.

**Results:** We built an age-of-onset axis for 2,885 Mendelian diseases from Orphanet and fitted cross-species conservation (241-way phyloP) and human-population constraint (*s*_het_) on the same 5,322 gene-disease pairs against matched controls, to test whether one decays faster with onset. Their onset slopes differed by 0.23 logit units (97.5% confidence interval −0.10 to 0.56), equivalent to 0.024 units of area under the receiver operating characteristic curve (AUROC) in a design resolving 0.049; the interval excludes constraint losing discriminative power faster than conservation by more than 0.011 AUROC units, widening to 0.033 when somatic associations are dropped. The usual explanation, that early-onset disease genes are developmental, pleiotropic and haploinsufficient by construction, did not account for the contrast; conditioning barely moved it. The answer did, however, depend on the constraint metric: three alternatives yielded larger contrasts excluding zero, though paired comparison could not distinguish the estimates. The metrics also diverged on recessive disease genes, with s_het_ ranked below the gene pool (AUROC 0.487) and loss-of-function depletion above it (0.533), though, against matched controls, both ranked them below (0.457 and 0.450). Evolutionary scores improved a non-evolutionary baseline by 17 to 23%, with no onset gradient.

**Conclusions:** Conservation and constraint do not lose their value at measurably different rates as disease onset rises, and with only 283 late-onset Mendelian diseases annotated genome-wide, the question cannot be settled more finely. The gene-class explanation does not account for the contrast. Metrics that rank genes near-identically can still yield opposite onset-stratified answers, so a result reported by one metric carries a sensitivity that its confidence interval does not show. No constraint metric identifies recessive disease genes against matched controls, so a pipeline with recessive disease in scope needs a correction that switching metric does not supply.

## Background

Gene-level evolutionary scores are among the most widely applied filters in human disease genetics. Constraint metrics derived from population sequencing rank candidate genes in exome and genome studies [1–3] and have been used for candidate prioritisation since the first large exome aggregations became available [4, 5]. Cross-species conservation supplies supporting evidence in variant classification frameworks [6, 7]. Genomic language models pretrained on evolutionary sequences are now proposed as an additional source of gene-level functional inference [8, 9]. Their common premise is that selection has recorded which genes matter, and it informs decisions beyond variant interpretation. Target-indication pairs with human genetic support reach approval roughly 2.6 times as often as those without, with the effect strengthening as confidence in the causal gene increases and remaining largely unaffected by genetic effect size, minor allele frequency and year of discovery [10]. The relationship was first quantified a decade ago [11, 12] and has since been extended to the majority of recently approved drugs [13]. Constraint therefore informs not only variant classification but also the choice of genes to pursue. The gnomAD analysis noted a limit to this premise: because reproduction is largely complete before late-life phenotypes appear, selection has little purchase on them, and a gene may tolerate inactivation on these measures despite causing severe or fatal late-onset disease [2]. This follows from classical theory: the force of selection declines with age once reproduction is substantially complete [14, 15], so deleterious alleles whose effects are confined to late life persist under weak purifying selection and may accumulate [16]. If the limit is sharp in practice, evolutionary evidence should become progressively less informative as disease onset rises, and least informative for the late-onset conditions that dominate clinical burden.

Several studies have tested this at the level of allele frequencies. Clustering 116 UK Biobank diseases into four age-of-onset groups found risk-allele frequencies varying systematically with the group’s onset profile, with support reported for both mutation accumulation and antagonistic pleiotropy [17]; an earlier analysis of disease-associated variants reached compatible conclusions [18]. Analysis of 62 diseases alongside longevity and fertility found evidence of positive selection on disease-risk alleles confined to conditions with onset below 40 years [19]. Against these, evolutionary-demographic modelling incorporating two-sex reproduction and grandmaternal care finds that purifying selection on late-onset susceptibility alleles is the norm rather than the exception [20, 21]. Whether the gradient appears in gene-level scores, which are the quantities used in practice, is a separate question. Constraint metrics summarise selection across all variants in a gene and are dominated by its core molecular function rather than by the fitness cost of any single phenotype. Evolutionary scores also differ from one another in the timescale over which they measure selection. Cross-species conservation [22, 23] integrates selection over approximately 100 million years across lineages whose life histories do not map onto human age of onset, and recent mammalian alignments have substantially increased the power to detect weak constraints without altering that property [24, 25]. Human-population constraint, whether expressed as a selection coefficient against heterozygotes [26, 27] or as depletion of observed loss-of-function variation [2, 3], reflects the recent human past and could, in principle, register onset-dependent fitness costs that phylogenetic averaging cannot. Two predictions follow, with opposite signs. If deep-time conservation is dominated by genes whose functions are required early in development, it should lose discriminative power more quickly as the onset rises. If instead human-population constraint registers the onset-dependent fitness costs that phylogenetic averaging cannot, constraint should be the score that decays, which is what the classical selection-shadow argument implies. The analysis plan registered the first, so a contrast above zero was predicted.

We therefore estimated a contrast rather than a level: the difference between the onset slopes of a conservation score and a constraint score, fitted jointly on the same genes and diseases. Disease-gene labels differ systematically in quality along the onset axis. Label noise attenuates both arms of a contrast, so the bias cancels in the difference to the extent that the two score families have equal main effects; both main effects are reported below with each family’s decay relative to its own main effect, as is the sensitivity of the contrast to the quality of the association labels. The study has three aims: to estimate this contrast across the full onset range of Mendelian disease, to test the gene-class explanation for the underlying constraint-onset relationship, and to characterise the dependence of the result on the choice of constraint metric. The analysis plan, filed in accordance with recommended practice for confirmatory analysis of existing data [28], is registered and publicly available [29].

## Methods

### Study design

This was a retrospective analysis of publicly available human genomic annotation. The unit of analysis was the gene-disease pair. The primary comparison was between two families of evolutionary score, cross-species conservation and human-population constraint, evaluated for their ability to discriminate disease genes from matched control genes as a function of the disease’s age of onset. The registered analysis plan [29] sets out the causal diagram, the two estimands, the assignment of scores to families, the onset binning, the adjustment set, the role of each covariate in it, the minimum detectable effect, and the stopping rules. Three elements of the fitted model differ from that plan. Correlation within disease and within gene is carried by a cluster-robust covariance instead of cross-classified random effects; onset enters linearly, with the midpoint age of each onset level as the sensitivity encoding, without the monotone spline the plan also named; and disease breadth, one of the two pleiotropy covariates the plan adjusts for, is excluded from every adjustment set because it counts the outcome among its inputs.

### Data sources

Mendelian disease-gene associations and age of onset were taken from Orphanet [30, 31] (en_product9_ages v1.3.42, released 23 June 2026; en_product6; en_product1) [32]. A gene-disease pair is entered as a positive when Orphanet records a disease-causing association, comprising germline mutations, those annotated as loss- or gain-of-function, and somatic mutations. Candidate-gene, biomarker, modifying, susceptibility-factor, fusion-gene and phenotype-role associations were not treated as positives. Dropping the somatic associations and the few not yet assessed is reported as a robustness arm. Complex disease associations, disease ontology, and molecular interactions were obtained from the Open Targets Platform release 26.06 [33]. Human-population constraint came from gnomAD v4.1, which provides LOEUF, pLI, and missense Z [2, 3], and from GeneBayes *s*_het_ [27], which extends the mutation-selection balance framework of earlier heterozygous selection coefficient estimates [26, 34] by incorporating gene features. Deep-time conservation was derived from UCSC phyloP [22], using the 241-way Zoonomia alignment as primary [24] and the 100-way alignment as secondary, and from phastCons [23]. Complex-disease onset was obtained from FinnGen release R13 via Risteys [35] and independently from the published median age at first record for 308 conditions in English National Health Service data [36]. Cell-line essentiality was the DepMap summary carried in the Open Targets target essentiality dataset [33, 37]. Tissue expression was the GTEx summary carried in the Open Targets baseline expression dataset [33, 38]. The Gene Ontology biological process annotation was used for the developmental gene-class covariate [39]. Trait-level pleiotropy was derived from published per-gene effect estimates across UK Biobank quantitative traits [40]. Balancing selection was summarised as standardised B_2_ aggregated per gene across three populations [41, 42]. The release of each source is given in Additional file 1, Table S1, and the per-gene conservation aggregates derived from these sources are given in Additional file 6.

### Onset axis

Mendelian onset used the earliest annotated Orphanet interval, on the reasoning that selection acts on the earliest fitness-relevant manifestation; the midpoint of each disease’s annotated interval span was used as the registered sensitivity encoding, which tests that rule rather than the spacing of the levels. Diseases annotated only as spanning every age carry no ordinal position and were excluded from the primary analysis, with their inclusion at the midpoint of that span as a named robustness arm. Seven Orphanet intervals were collapsed to five under a rule that merged levels with fewer than 50 diseases into the adjacent level, applied before any score was examined. Complex-disease onset was the median age at first recorded event, binned by decade, after a hundred-case floor. Registry endpoints were mapped to disease ontology terms via their EFO annotations, and obsolete identifiers were resolved to their Mondo replacements. Mappings whose endpoint and term names disagreed were adjudicated by the relationship between the two terms in the Mondo hierarchy, and these 428 mappings, with route, name agreement and verdict, are given in Additional file 5.

### Statistical model

For gene *g* and disease *d*, with *y_g,d_* = 1 where *g* is a positive for *d*:

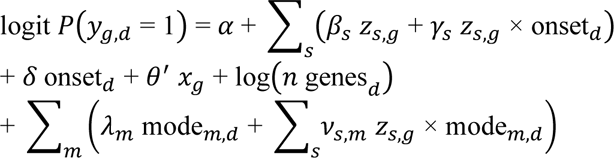

where *z_s,g_* is the score s standardised within the analysis set, onset is the disease’s onset level scaled to the unit interval and centred, and *x_g_* comprises coding sequence length decile, expected loss-of-function count decile and GC content. The index s runs over the two score families, conservation and constraint, and m over the two resolved inheritance modes; the terms mode*_m,d_* are indicators for dominant and for recessive inheritance, with mixed or unknown mode as the reference. α is the intercept, *β_s_* and δ the main effects of score and of onset, *γ_s_* the score-by-onset interaction whose contrast defines Γ below, θ the vector of coefficients on *x_g_*, *λ_m_* the main effect of inheritance mode, and *ν_s,m_* the score-by-mode interaction. The log count of genes annotated to disease d enters without a fitted coefficient. Every score is oriented so that a higher value means more constrained or more conserved; LOEUF, an upper bound on the ratio of observed to expected loss-of-function variants, is negated before standardisation. In the conditioning specifications, each gene-class covariate is standardised and enters as a main effect together with its interaction with each score, so a covariate capable of carrying the score-by-onset interaction is given the terms that would absorb it. Rows missing any covariate in the block are dropped rather than imputed, and the contrast on all rows is reported beside every conditioned estimate.

Genes recur across diseases, and diseases cluster their genes, so rows are correlated in two directions, and the contrast needs a standard error that accounts for both. The model was fitted as a logistic regression with a two-way cluster-robust sandwich covariance estimator, following Cameron, Gelbach, and Miller [43]. It is consistent under arbitrary correlation within a disease and within a gene, assumes no distribution for either, and is fast enough to carry a permutation null of several thousand replicates. Its coefficients are population-averaged, which is what a contrast between two interaction terms calls for. The design-stage power simulation clustered on disease alone, so the minimum detectable effects reported here are computed from the realised standard error of each fitted contrast instead. Each evidence source was fitted on its own rows, with no pooled source term, because label quality and pleiotropy differ across sources. Onset entered linearly, with the midpoint of each disease’s annotated interval span as the registered sensitivity encoding. No minimum gene count filter was applied to the primary contrast. A ten-positive threshold applies to the incremental-value estimand, which requires a per-disease discrimination estimate, and to per-disease points drawn on figures. The primary estimand is Γ = γ_constraint_ − γ_conservation_, the difference between the two score-by-onset interaction coefficients, taken as a contrast on the fitted covariance. Because a score’s decay along onset is the negative of its interaction coefficient, Γ above zero means conservation loses discriminative power faster than constraint as onset rises. Controls were matched ten per positive on confounders only, without replacement within disease. Both matched and unmatched analyses are reported. Every row of the fitted design, with its gene, disease, onset position, positive status and model terms, is provided in Additional file 3. Mode of inheritance is entered as a stratifying interaction with each score because s_het_ is a selection coefficient on heterozygotes [26, 27], and the dominant share of Mendelian diseases rises along the onset axis. Gene-level discrimination within each inheritance stratum was measured against two reference sets: the pool of scored genes, and ten controls per disease gene drawn from genes causing no Mendelian disease that share the gene’s stratum of coding sequence length decile, expected loss-of-function count decile and GC decile. Controls were drawn without replacement within a gene and could serve more than one gene, as they are drawn within a disease in the primary design. Intervals for the matched comparison resample the matched blocks, each disease gene carried with its own controls. Both comparisons are exploratory, and neither was registered.

### Incremental value over a non-evolutionary baseline

For each disease, fivefold cross-validated discrimination was computed for nested models: M0 with non-evolutionary features only, comprising expression, tissue specificity, network topology, and cell-line essentiality, and M1 adding evolutionary scores. The estimand is the slope on the onset of log *R*, where *R* = (AUC_M1_ − 0.5) / (AUC_M0_ − 0.5), restricted to diseases with AUC_M0_ − 0.5 > 0.05. The ratio form is invariant to multiplicative attenuation from label noise, which varies along the onset axis.

### Power

Power was simulated for the realised design using the disease- and gene-disease-pair counts, the 10 matched controls per positive, and α = 0.025 for the fitted model. Each cell of the contrast grid was run with 1,000 replicates, and each cell of the incremental-value grid with 800. The main effect was defined as the pooled marginal discrimination of the constraint score against matched controls, with no score-by-onset interaction. That simulation clustered standard errors on disease alone and returned 0.035 AUROC for the primary contrast, 0.034 with random controls, 0.041 with two onset levels, and 0.052 when restricted to dominant disease (Additional file 1, Section 4). The minimum detectable effects reported here are computed as the realised standard error of each fitted contrast multiplied by 3.08, the sum of the normal quantiles at α = 0.025, two-sided, and 80% power. The realised standard error is 0.147, compared with 0.106 implied by the simulation, a factor of 1.17 due to the second clustering dimension. A minimum detectable effect computed this way uses the design’s precision rather than the observed effect, which is what reporting it alongside a null estimate requires [44]. The incremental-value simulation assigned each disease its own measured baseline discrimination and its own count of positives, with the sampling error for the per-disease AUROC estimated from the Hanley-McNeil expression [45]. Its grid was indexed by the true ratio and by the correlation between the nested AUROCs. That correlation cannot be identified from the data, so the simulated grid is read at a ratio of 1.3 and zero correlation, the most conservative of the correlations simulated, which gives 0.70 and 0.47. The minimum detectable slopes reported in Results are computed from the realised standard error of each fitted slope, on the same rule as the contrast.

### Inference

Intervals on Γ and its components, on the control decays and on the incremental-value slope are Wald intervals from the two-way cluster-robust covariance at α = 0.025, the error rate the design was powered at, and so have 97.5% coverage. All other intervals are 95% bootstrap percentile intervals, based on 400 resamples of diseases for paired differences, movements under conditioning, and correlations with onset, and on resamples of genes for discrimination. Differences between quantities were tested as paired differences within a common design, rather than by comparing marginal intervals, since the variance of a difference between estimates fitted to shared data depends on their correlation, which marginal intervals do not capture [46]. Each adjustment covariate was characterised by its gradient with onset and by whether the outcome appears among its inputs. The permutation null used 8,000 draws with onset permuted within the therapeutic area, enough that the Monte Carlo error on the null’s centre is small against the displacement being measured.

### Genomic language model scoring

Evo 2 was scored with the hosted 40B model, the size used to compute the published benchmark [9]. The insertion rule was an 8-kb window with stop codons at nucleotide offsets of 15, 30, and 45. Inclusion in the conservation family required discriminating DepMap essential genes [37] within 0.03 of the published AUROC of 0.66 [9], which was obtained using a different perturbation (Additional file 1, Section 9).

### Use of large language models

Artificial intelligence was used during the preparation of this manuscript. Claude (Anthropic; models Claude Opus 5 and Claude Sonnet 5) was used for language editing of author-written text and for coding assistance during implementation of the analysis pipeline. No text, figures, or data were generated de novo by the tool, and all AI-assisted output was reviewed, verified, and edited by the authors, who take full responsibility for the content, accuracy, and integrity of this study.

## Results

### An age-of-onset axis for Mendelian disease

Age of onset was taken from the Orphanet natural history product [30, 32], which annotates 6,982 of 7,790 ORPHAcodes. Excluding 868 annotated only as spanning all ages, 6,114 carry an ordinal interval, and 3,524 carry both an ordinal interval and a gene association (Fig. 1a). Seven Orphanet intervals were collapsed to five levels, merging antenatal with neonatal and elderly with adult (Additional file 1, Section 1). The Mendelian arm spans all levels, with at least 100 diseases and 200 gene-disease pairs (Table 1, Fig. 1b), and the onset, inheritance, and mean score for each fitted disease are listed in Additional file 4. Complex-disease sources are almost entirely unassigned on this axis, as expected given that Orphanet annotates rare disease [31], and were placed on a separate axis derived from the FinnGen registry [35]. The adult level contains 283 diseases and 484 gene-disease pairs, and its composition is not a random draw from the genome: 51 are neurodegenerative, 49 are hereditary cancer predisposition, 29 are metabolic, 26 are cardiac or neuromuscular, and 128 fall outside these categories. Dropping hereditary cancer predisposition, whose genes are DNA repair genes constrained independently of when cancer presents, moved the primary estimate to 0.22. The dominant share of Mendelian diseases rises from 0.32 at perinatal onset to 0.71 at adult onset (Fig. 1c). Because *s*_het_ is a selection coefficient on heterozygotes and discriminates dominant from recessive inheritance [26], this shift would by itself raise the apparent discrimination of constraint at later onset. Median age at first recorded event in FinnGen agreed with an independent English registry series [36] at Spearman 0.867 across 161 matched conditions, with FinnGen later by a median of 2.3 years (Fig. 1d). Left truncation was quantified against the same series: for conditions with an English median below 40 years, the Finnish registry records first events 10.2 years later on average, against 1.3 years at 40 and above (Additional file 1, Section 3).

**Fig. 1.**
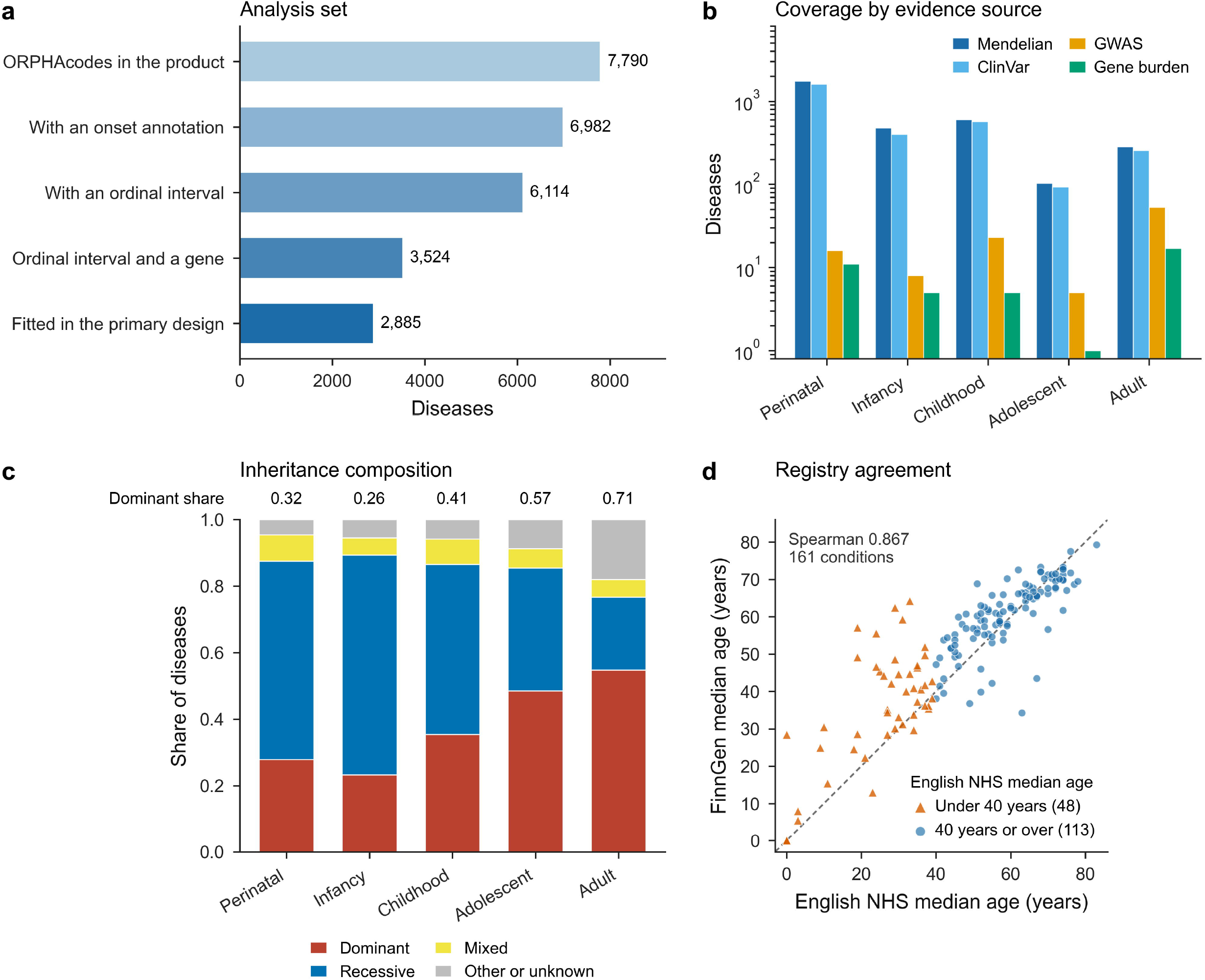
Age-of-onset axis and composition of the analysis set. **a** Derivation of the analysis set from Orphanet annotations, from 7,790 ORPHAcodes to the 2,885 diseases fitted in the primary design. The gene step counts any association in the Orphanet gene product, a broader set than the gene associations that enter the analysis, which is why the Mendelian onset levels of Table 1 sum to 3,213 rather than 3,524. **b** Diseases by onset level and evidence source, on a logarithmic scale. **c** Inheritance composition by onset level, with the dominant share of diseases carrying a resolved mode above each bar, rising from 0.32 at perinatal onset to 0.71 at adult onset. **d** Median age at first recorded event in FinnGen against the English National Health Service series for 161 matched conditions, with the identity line. Conditions with an English median below 40 years, drawn as triangles, carry a left-truncation offset of 10.2 years, compared with 1.3 years above that age.

**Table 1.** Diseases and gene-disease pairs by onset level and evidence source.

| Onset level | Mendelian diseases | Mendelian pairs | GWAS diseases | ClinVar diseases | Gene burden diseases |
| --- | --- | --- | --- | --- | --- |
| Perinatal | 1,749 | 3,263 | 16 | 1,610 | 11 |
| Infancy | 477 | 851 | 8 | 399 | 5 |
| Childhood | 601 | 1,124 | 23 | 568 | 5 |
| Adolescent | 103 | 209 | 5 | 93 | 1 |
| Adult | 283 | 484 | 53 | 257 | 17 |
| Unassigned | 616 | 1,098 | 9,747 | 6,569 | 1,201 |

Counts are of diseases carrying an onset annotation and at least one gene association in each evidence source, before the restrictions that define the fitted design. The Mendelian design additionally requires an autosomal gene carrying both primary scores, which leaves 2,885 diseases and 5,322 gene-disease pairs. Perinatal merges the antenatal and neonatal Orphanet intervals; adult merges the adult and elderly intervals. The adult Mendelian cell contains 128 diseases outside the named categories: 51 neurodegenerative, 49 hereditary cancer predisposition, 29 metabolic, and 26 cardiac or neuromuscular.

### Conservation and constraint decay at indistinguishable rates

The contrast between the onset slopes of 241-way phyloP and *s*_het_ was 0.23 in logit units (97.5% CI −0.10 to 0.56), equivalent to 0.024 AUROC units (Table 2, Fig. 2a). The minimum detectable effect of the realised design is 0.049 AUROC units, or 0.45 logit units, at 80% power and α = 0.025. Read as a one-sided bound, the interval excludes constraint losing discriminative power faster than conservation, the direction the classical selection-shadow argument predicts, by more than 0.102 logit units, or 0.011 AUROC units. With the somatic and not-yet-assessed associations dropped, the same bound is 0.309 logit units, or 0.033 AUROC units. A permutation null with onset shuffled within therapeutic area over 8,000 draws gave a two-sided *P* of 0.099 (Fig. 2b). Model-implied discrimination fell across the onset axis at a similar rate in both families (Fig. 2c). The contrast decomposes into a conservation decay of 0.360 (97.5% CI 0.180 to 0.539) and a constraint decay of 0.132 (−0.061 to 0.326), against main effects of 0.263 (0.120 to 0.406) and 0.212 (0.005 to 0.420). Relative to its own main effect conservation decays at 1.37 and constraint at 0.62, and the conditioned and label-restricted fits are in Additional file 1, Table S16. In the control analyses, neither negative control carried an onset gradient (Table 3, Fig. 2d). Developmental annotation, the positive control, decayed at 0.15 (97.5% CI 0.04 to 0.26). The permutation null over 8,000 draws was centred at −0.009, displaced from zero by 0.06 of its own standard deviation, with a central 95% span of −0.28 to 0.25, whose implied standard deviation of 0.136 is close to the 0.147 standard error of the fitted contrast.

**Fig. 2.**
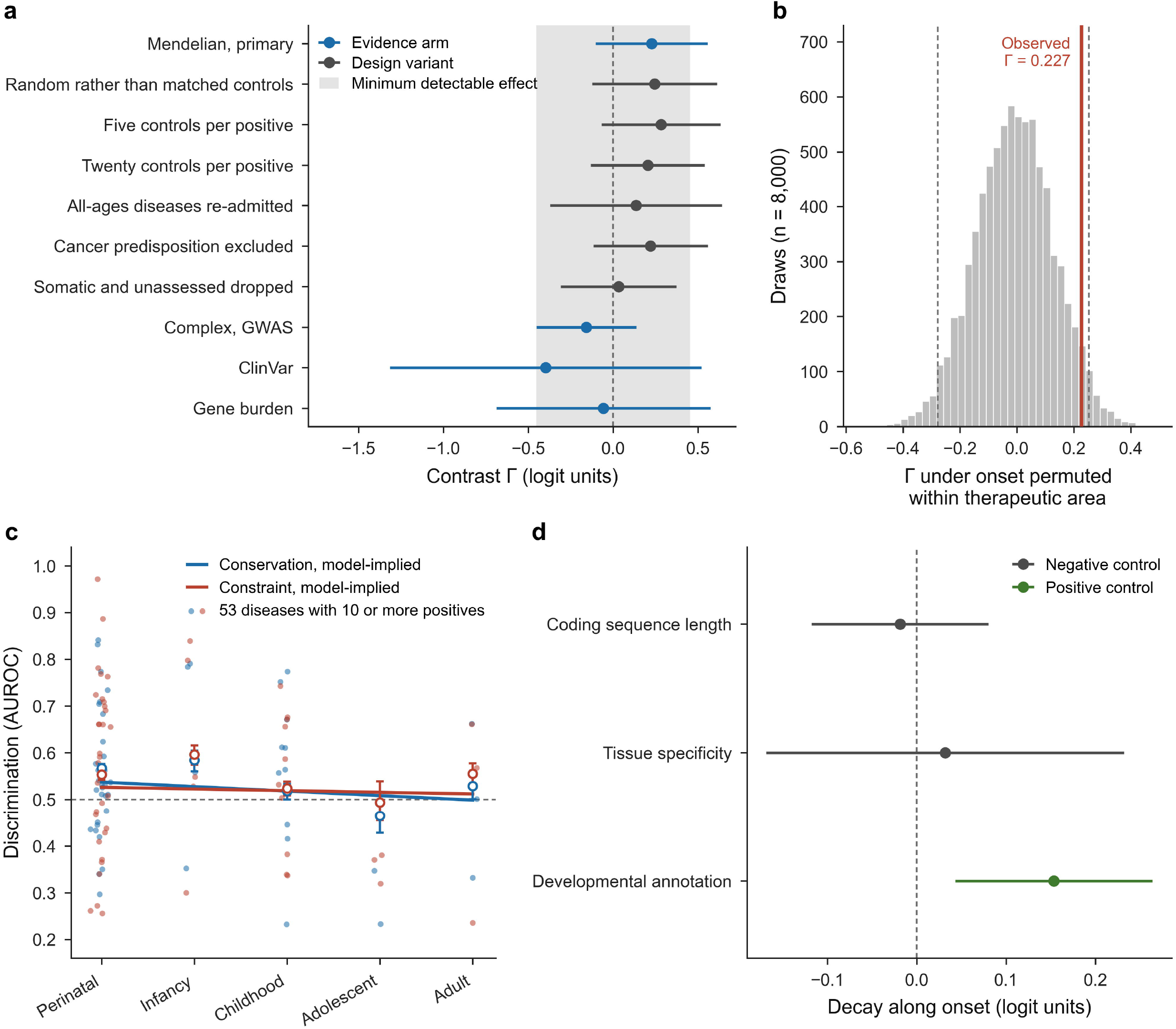
The onset contrast between conservation and constraint. **a** Contrast estimates with 97.5% intervals across evidence arms and design variants. The shaded band marks the minimum detectable effect of the realised Mendelian design, 0.45 logit units either side of zero. **b** Permutation null over 8,000 draws with onset shuffled within therapeutic area, with the central 95% span dashed and the observed estimate marked. **c** Model-implied discrimination across the five onset levels for each score family, drawn as a line, with the discrimination measured within each level as open circles and 95% intervals, and per-disease values for the 53 diseases having at least ten positives as small points. **d** Control analyses with 97.5% intervals, showing the two negative controls and the positive control.

**Table 2.**
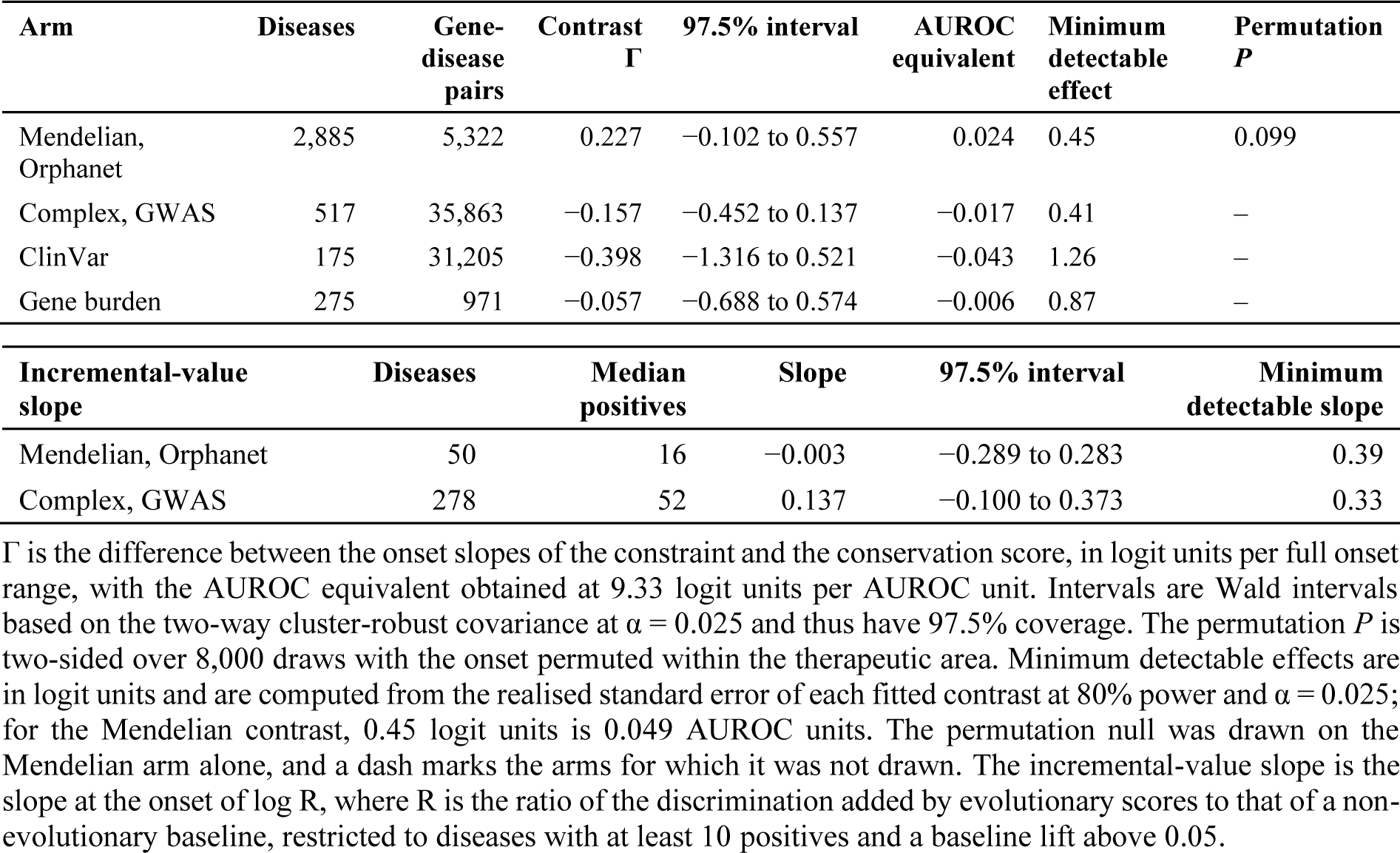
Contrast estimates and incremental-value slopes with minimum detectable effects.

**Table 3.**
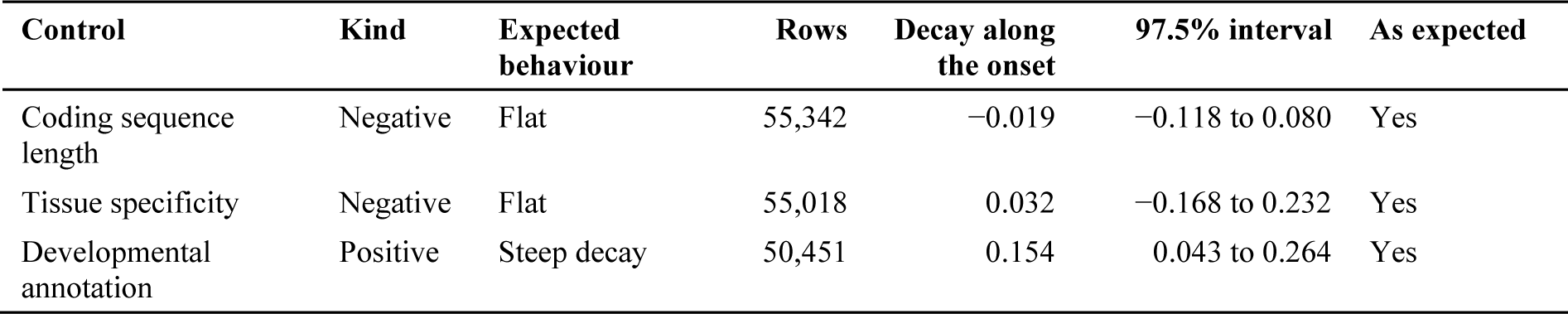
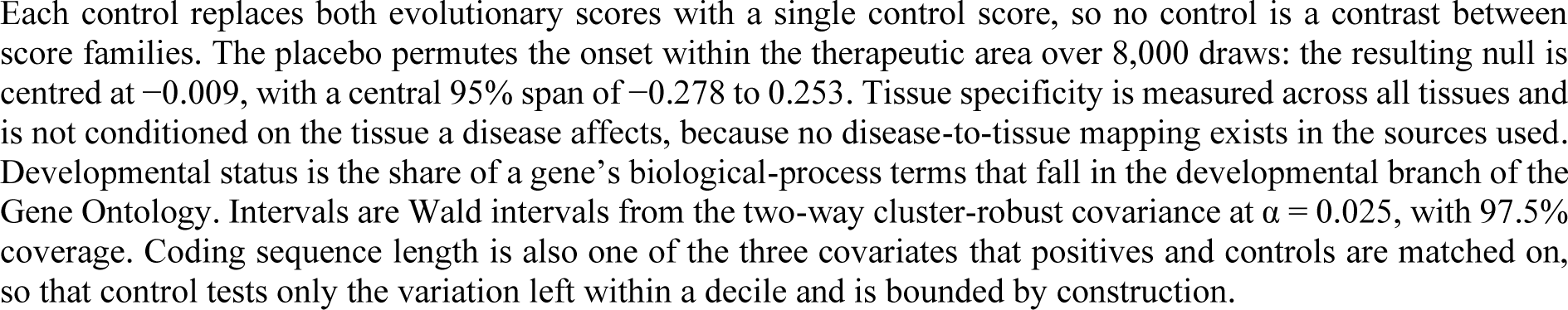
Control analyses with expected and observed behaviour.

The estimate was stable across most design variants (Table 4, Fig. 2a). Removing the genes-per-disease term moved it from 0.2275 to 0.2269. Random rather than matched controls gave 0.245, and five and twenty controls per positive gave 0.283 and 0.204. Re-admitting the diseases annotated only as spanning every age, at the midpoint of that span, gave 0.135 (97.5% CI −0.371 to 0.642) on 3,094 diseases and 5,750 gene-disease pairs. Coding onset as the midpoint of each disease’s annotated span gave 0.163 (97.5% CI −0.167 to 0.492). Dropping the 244 Orphanet associations recording somatic mutations and the 17 not yet assessed, which are concentrated at adult onset, moved the contrast to 0.032 (97.5% CI −0.309 to 0.373); across five matching seeds, the contrast ran 0.179 to 0.236 on the full set and 0.032 to 0.124 on the restricted one, so the movement is about twice the variation the control draw alone produces. In the complex-disease arm, the contrast was −0.16 (97.5% CI, −0.45 to 0.14; Table 2), and neither the ClinVar nor the gene-burden arm reached significance. The Mendelian and complex point estimates have opposite signs but overlapping intervals from −0.10 to 0.14, and cover different onset ranges: antenatal to adult versus approximately 25 to 75 years.

**Table 4.** Contrast estimates across constraint and conservation metrics with paired differences.

| Specification | Gene-disease pairs | Contrast $\Gamma$ | 97.5% interval | Standard error | Excludes zero |
| --- | --- | --- | --- | --- | --- |
| $S_{het}$, primary metric | 5,322 | 0.227 | $-0.102$ to $0.557$ | 0.147 | No |
| LOEUF, gnomAD v4.1 | 5,360 | 0.466 | $0.280$ to $0.652$ | 0.083 | Yes |
| Missense Z | 5,360 | 0.377 | $0.133$ to $0.621$ | 0.109 | Yes |
| LOEUF, gnomAD v2.1.1 | 5,320 | 0.475 | $0.308$ to $0.643$ | 0.075 | Yes |
| phyloP 100-way, conservation swap | 5,322 | 0.183 | $-0.132$ to $0.499$ | 0.141 | No |
| phastCons, conservation swap | 5,322 | 0.116 | $-0.186$ to $0.419$ | 0.135 | No |
| phyloP fourfold degenerate, conservation swap | 5,322 | $-0.001$ | $-0.231$ to $0.230$ | 0.103 | No |
| Onset slope by inheritance mode, full sample | 5,322 | 0.429 | $-0.220$ to $1.077$ | 0.289 | No |
| All-ages diseases re-admitted | 5,750 | 0.135 | $-0.371$ to $0.642$ | 0.226 | No |
| Somatic and unassessed associations dropped | 5,155 | 0.032 | $-0.309$ to $0.373$ | 0.152 | No |
| Onset coded as the midpoint of each disease's annotated span | 5,322 | 0.163 | $-0.167$ to $0.492$ | 0.147 | No |

| Paired difference on a common gene set | Difference | 95% interval | Resamples |
| --- | --- | --- | --- |
| LOEUF minus $S_{het}$ | 0.279 | $-0.037$ to $0.551$ | 400 |
| Missense Z minus $S_{het}$ | 0.174 | $-0.017$ to $0.391$ | 400 |

Each row swaps one score for another while leaving the rest of the design unchanged, so the pair count varies with the metric's coverage. Contrast intervals are Wald intervals from the two-way cluster-robust covariance at $\alpha = 0.025$ , with 97.5% coverage. Paired differences are fitted to the genes that carry all three constraint metrics and resampled using the same disease draws, with 95% bootstrap percentile intervals. Standard errors are those of the fitted contrast in each arm.

### Gene class does not account for the contrast

Constraint is expected to track the onset because early-onset disease genes are developmental, pleiotropic, and haploinsufficient by construction, so that the relationship may reflect gene class rather than the fitness cost of disease. This account requires both that these properties co-vary with onset among disease genes and that conditioning on them removes the contrast, and the data support neither. Conditioning bounds confounding of the contrast rather than of either score’s onset slope, so an account under which gene class steepens both slopes equally is not addressed by this evidence. Only one of the three properties varies with onset among disease genes (Fig. 3a). The share of developmental annotation has a rank correlation with onset of −0.083 (95% CI, −0.121 to −0.048). Haploinsufficiency does not vary (0.014, 95% CI −0.034 to 0.062), and neither does trait-level pleiotropy (0.011, 95% CI −0.015 to 0.037). Inheritance composition runs counter to the account, with the dominant share rising rather than falling across the onset. Conditioning on developmental annotation did not attenuate the contrast under any constraint metric (0.007 for *s*_het_, −0.013 for LOEUF and 0.002 for missense Z; all intervals containing zero). Across the three retained covariates and three metrics, the lowest interval bound was −0.024, compared with contrasts ranging from 0.23 to 0.50 (Fig. 3b; Additional file 1, Section 7). Two of the nine cells carried intervals excluding zero, both under haploinsufficiency (LOEUF 0.035, 95% CI 0.021 to 0.055; missense Z 0.024, 95% CI 0.013 to 0.045). pLI and LOEUF derive from the same gnomAD loss-of-function counts [2], so conditioning one on the other is collinear rather than independent, and neither movement supports the converse claim that gene class suppresses the relationship.

**Fig. 3.**
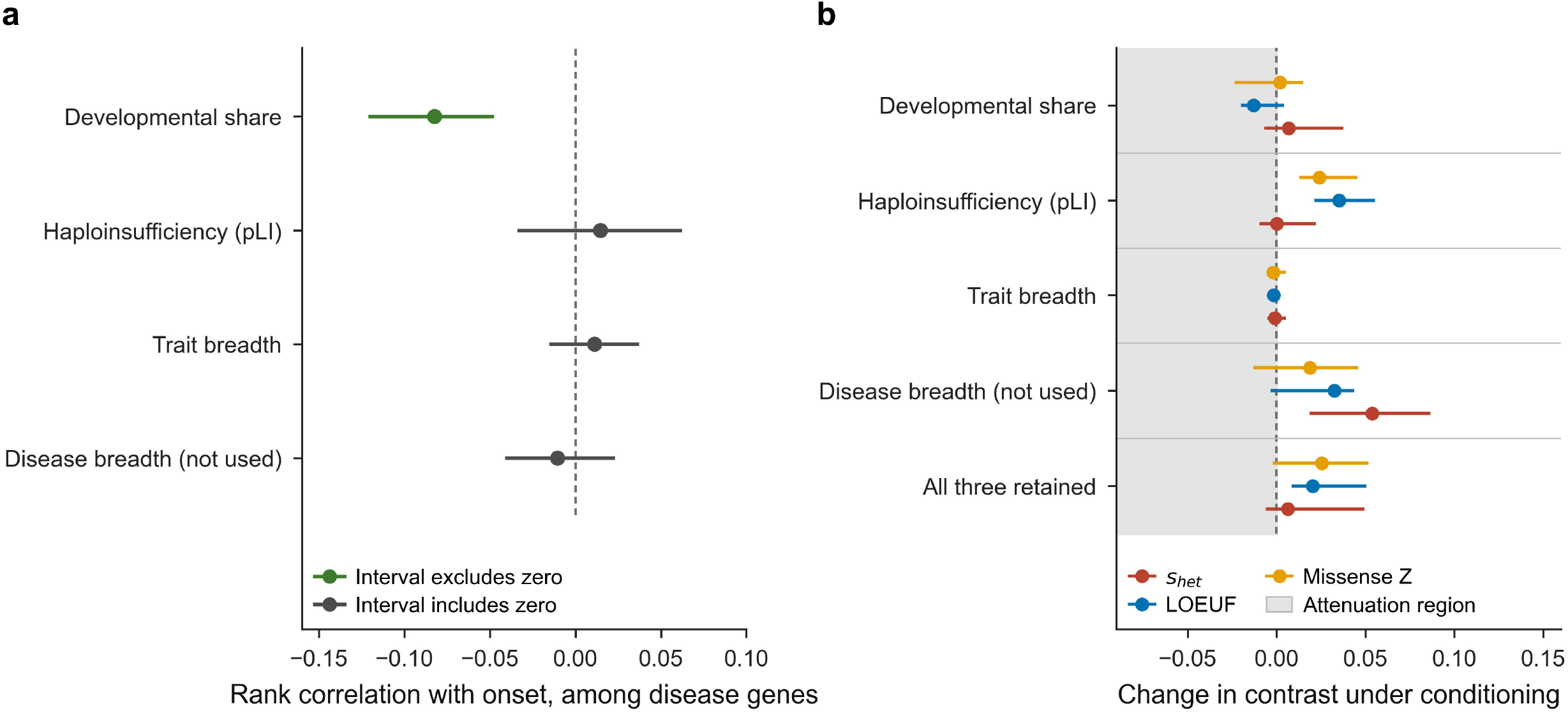
Gene class and the onset contrast. **a** Rank correlation of each gene-class covariate with onset among disease genes, with 95% intervals. Only the developmental share varies; disease breadth is shown for completeness and is not used as an adjustment covariate. **b** Change in the contrast under conditioning, for each covariate and constraint metric and for the three retained covariates together, with 95% intervals. The shaded region is where a change would attenuate the contrast. Cells whose covariate does not vary with onset in panel a are uninformative, not supportive.

### An annotation-derived pleiotropy measure encodes the outcome

Two measures of pleiotropy were included in the adjustment set. Trait breadth, the effective number of traits from per-gene effect estimates across UK Biobank quantitative traits [40], moved the contrast by less than 0.002 under every metric. Disease breadth, a count of therapeutic areas from Open Targets evidence [33], carried the entire movement on the primary metric (0.054, 95% CI 0.019 to 0.087). Disease breadth is assembled from evidence sets that include Orphanet associations that supply the positives, so it counts the outcome among its inputs. It is therefore excluded from every adjustment set and reported separately. The behaviour is an instance of multifunctionality bias, in which apparent prediction performance derives from gene annotation depth rather than from the associations being modelled [47–49]. Developmental annotation share correlates with a gene’s biological-process term count at 0.296 here, comparable to the 0.34 correlation with publication count reported for autism prioritisation scores [48]. The underlying asymmetry in annotation depth is well characterised: a minority of genes accumulate most of the literature, and the reasons are largely historical rather than biological [50]. The study by Gillis and Pavlidis left open whether highly ranked genes are genuinely more multifunctional or appear so through annotation bias [47]. Comparing an assay-derived and an annotation-derived measure on the same genes and model favours annotation bias, although trait breadth and disease breadth do not measure one construct, and a covariate drawing on the evidence that defines the outcome would move the estimate whatever pleiotropy did.

### Constraint metrics that agree on gene ranking disagree on the onset contrast

Three alternative constraint metrics were examined. All three gave larger contrasts than *s*_het_ and all three excluded zero (Table 4): LOEUF from gnomAD v4.1 at 0.47 (97.5% CI 0.28 to 0.65), missense Z at 0.38 (0.13 to 0.62), and LOEUF from gnomAD v2.1.1 [1, 2] at 0.48 (0.31 to 0.64). Conservation swaps behaved differently, with 100-way phyloP at 0.18 and phastCons at 0.12, both null. Aggregating phyloP at fourfold degenerate sites, where coding constraint is weakest, and the score carries the least information about the gene, returned −0.001 (−0.231 to 0.230), the arm closest to a pure negative control among the score swaps. Decomposing the contrast locates the point-estimate pattern in the constraint component (Fig. 4a). With conservation held constant, its decay runs from 0.35 to 0.41 across specifications, while the constraint decay changes sign: 0.144 under *s*_het_, 0.039 under missense Z and −0.078 under LOEUF. The metrics disagree on the direction of their own onset slopes while ranking genes similarly at every onset level (Spearman −0.755 to −0.878 for *s*_het_ against LOEUF, and −0.821 across all scored genes; Fig. 4b). The arms also differ in precision, in the direction predicted by their score correlations. Across scored genes, the primary conservation aggregate correlates with s_het_ at 0.613, LOEUF at 0.525, missense Z at 0.409, and the fourfold degenerate aggregate at 0.334. The standard error of a difference between two standardised coefficients rises as the two regressors converge, and the arms order accordingly: 0.147 logit units for the primary pairing, 0.141 and 0.135 for the conservation swaps that retain *s*_het_, 0.103 for the fourfold swap, and 0.075 to 0.109 for the three arms that replace *s*_het_. Part of the metric dependence is therefore a difference in precision rather than in what the metrics measure, since *s*_het_ is estimated from gene features that include cross-species conservation [27], which leaves the primary contrast setting conservation against a score partly derived from it. The correlation accounts for most of the gap to the fourfold swap and for about a fifth of the gap to LOEUF.

**Fig. 4.**
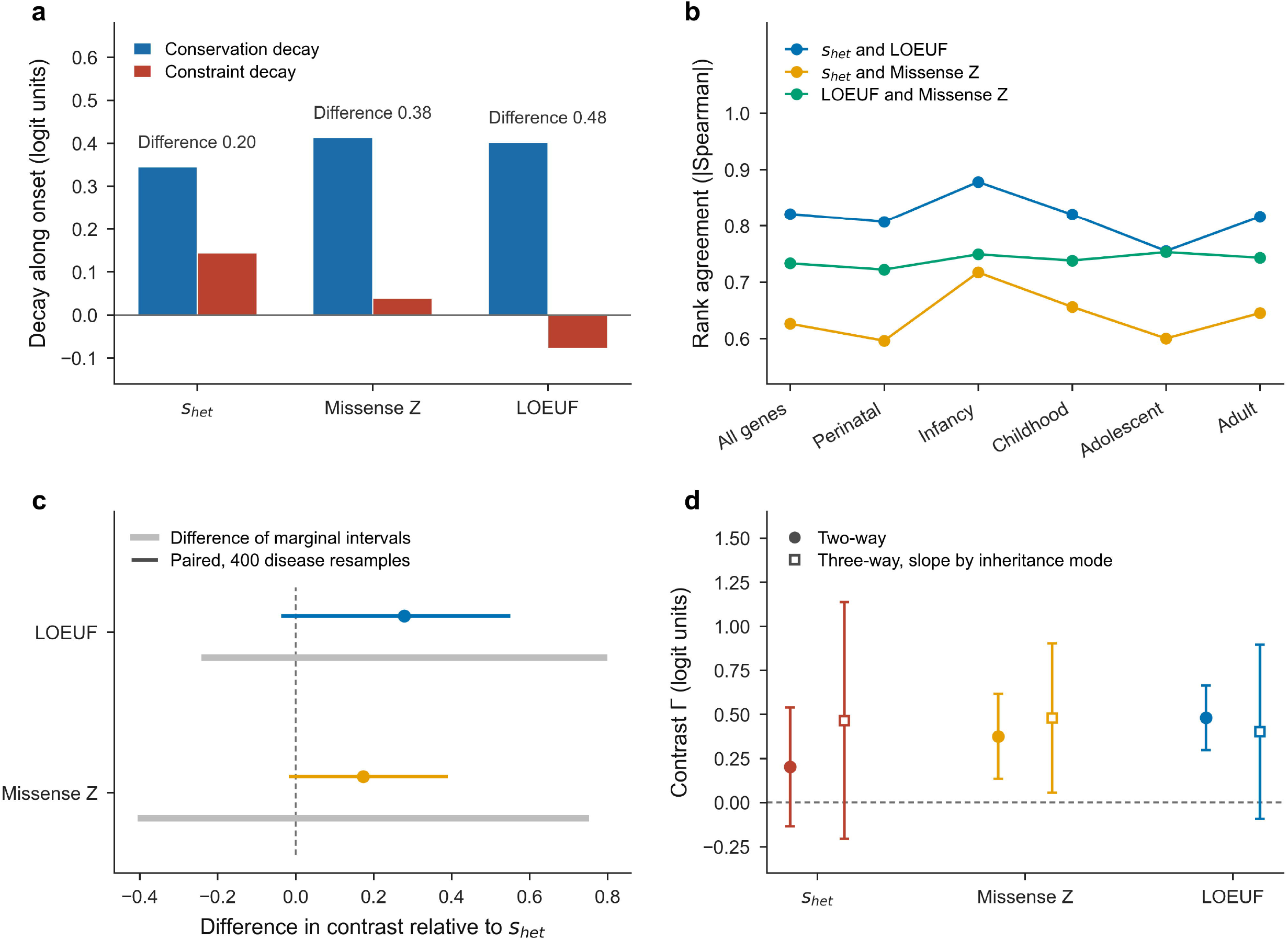
Dependence of the contrast on the choice of constraint metric. **a** Conservation and constraint components of the contrast under each constraint metric, fitted on the genes carrying all three metrics, with the difference between the two components above each pair. The constraint component changes sign across metrics. **b** Rank agreement among constraint metrics within each onset level, as the absolute Spearman correlation. **c** Paired differences between metrics, bootstrapped over 400 resamples of diseases, with the difference between the marginal intervals from the same fit drawn behind each in grey. **d** Contrast estimates for each constraint metric under the two-way and the three-way inheritance specification, with 97.5% intervals. Both are fitted on the genes carrying all three metrics, so the panel compares specifications rather than samples; the three-way estimate on the full sample is in Table 4.

Paired comparisons of the genes carrying all three metrics showed that the estimates are not separable. LOEUF minus *s*_het_ was 0.28 (95% CI −0.04 to 0.55), and missense Z minus *s*_het_ was 0.17 (−0.02 to 0.39), bootstrapped over 400 resamples of diseases (Table 4, Fig. 4c). The metric chosen changes the conclusion, although the underlying estimates are not shown to differ. We tested three mechanisms, and none accounted for the pattern. If a feature-informed estimator [27] carries gene-class information, conditioning on gene class should recover more contrast for *s*_het_ than for direct measures; *s*_het_ moved least. If inheritance composition drove the pattern, the dominant-to-recessive gap should be wider for LOEUF than for s_het_; it is narrower, 0.19 against 0.24, and 0.20 against 0.22 on matched controls. Allowing the onset slope to vary by inheritance mode converges the point estimates, with spread across metrics falling from 0.28 to 0.08 on the common gene set, but widens intervals to between 0.85 and 1.34, so the metrics cannot be distinguished from each other or, in two of three cases, from zero (Fig. 4d). Two further quantities bear on the gap. The first is the movement under conditioning, 0.006 to 0.027, an order of magnitude below the 0.28 point-estimate gap, which rules out covariate confounding of comparable size. The second is the correlation between the two contrast estimates across the 400 disease resamples. It is −0.14 for LOEUF and 0.59 for missense Z against *s*_het_, both below the 0.82 and 0.63 thresholds at which the same scores rank genes, so the sampling error of each contrast depends on which diseases are resampled and is largely metric-specific.

### s_het_ and depletion measures diverge on recessive disease genes, but not against matched controls

Discrimination was measured for 2,173 recessive and 1,218 dominant disease genes against a pool of 17,434 scored genes, with 1,000 bootstrap resamples (Fig. 5a). On recessive disease genes *s*_het_ discriminated at AUROC 0.487 (95% CI 0.476 to 0.498), below chance. LOEUF and missense Z were above chance on the same genes at 0.533 (0.521 to 0.544) and 0.521 (0.509 to 0.534), respectively. Paired on the same genes, LOEUF exceeded *s*_het_ by 0.046 (95% CI 0.037 to 0.055; Fig. 5a). That comparison is made against the unmatched pool. Coding sequence length and expected loss-of-function count, two of the covariates the primary design matches on, separate recessive disease genes from the pool on their own, and LOEUF is a function of the expected count. Repeating the comparison against 10 controls per gene, matched on those covariates, removes the difference and reverses its sign. *s*_het_ reached 0.457 (95% CI 0.447 to 0.469), LOEUF 0.450 (0.441 to 0.460) and missense Z 0.472 (0.462 to 0.483), so LOEUF minus *s*_het_ was −0.007 (−0.014 to −0.0004) and missense Z minus *s*_het_ was 0.015 (0.005 to 0.025). All three sit below chance, so no constraint metric identifies recessive disease genes among genes of the same coding length, expected loss-of-function count and base composition. For dominant disease genes, the matched comparison reverses the metrics, with *s*_het_ at 0.674 (0.657 to 0.690) above LOEUF at 0.650 (0.634 to 0.664) and missense Z at 0.634 (0.618 to 0.647). Matching left standardised mean differences of 0.07 on coding length, 0.10 on expected loss-of-function count and 0.01 on GC content, against 0.31, 0.40 and −0.04 unmatched, and supplied 9.6 controls per gene, with 154 of the 2,173 recessive genes in strata holding fewer than ten (Additional file 1, Section 8). Two readings are possible. *s*_het_ may be underperforming, since a score blind to recessive genes would sit at 0.5 rather than below it. Alternatively, it may be behaving as defined, since *s*_het_ estimates selection on heterozygotes [26, 34] and recessive disease genes have unaffected heterozygotes, in which case LOEUF at 0.533 against the pool is the value requiring explanation, and the matched comparison supplies it. The original formulation reported that a threshold on *s*_het_ discriminates dominant from recessive inheritance with high positive predictive value [26], which is the same property viewed in reverse. The displacement from chance is small in either case, at 0.013 against the pool and 0.043 against matched controls, and resolved only because 2,173 recessive disease genes contribute. For prioritisation, the consequence is the same under both readings: *s*_het_ ranks recessive disease genes below the genome average. Loss-of-function depletion measures rank them above it, but they do so by coding length and expected loss-of-function count rather than by constraint, and, against controls holding those fixed, they rank them below it as well.

**Fig. 5.**
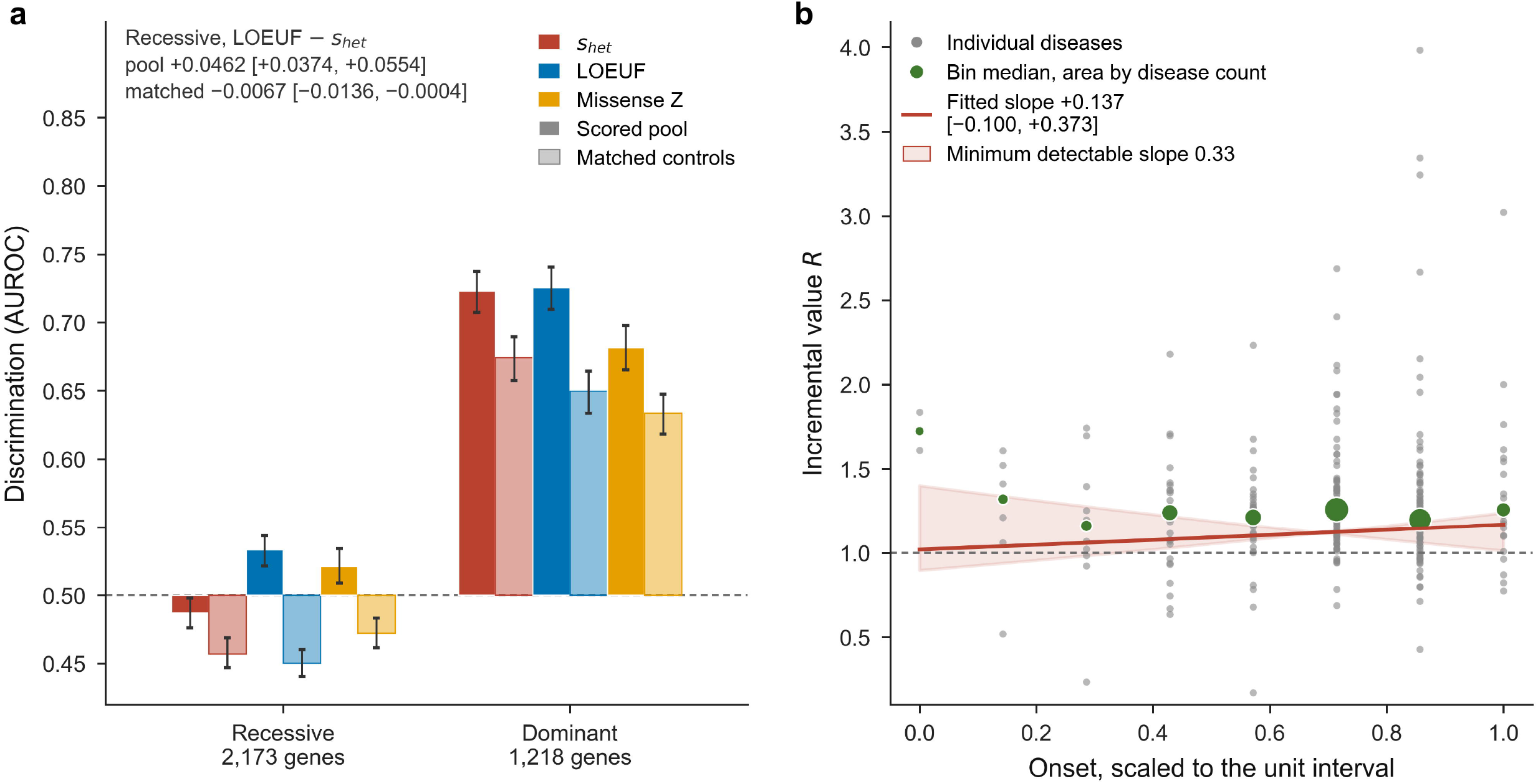
Divergence on recessive genes and incremental value across onset. **a** Discrimination by inheritance stratum and constraint metric against two reference sets, the pool of 17,434 scored genes in full colour and ten controls per gene matched on coding sequence length, expected loss-of-function count and GC content in pale, with 95% intervals and the paired difference between LOEUF and *s*het on recessive genes annotated for both. The chance line is marked at 0.5. **b** Incremental-value ratio against onset on the complex-disease arm, with individual diseases in grey, bin medians sized by disease count, the fitted slope, and the band the minimum detectable slope of 0.33 would span.

### Evolutionary scores add about a fifth to a non-evolutionary baseline, with no detectable onset gradient

Adding evolutionary scores to a model built from expression [38], tissue specificity, network topology [33] and cell-line essentiality [37] raised discrimination by a median factor of 1.17 on the Mendelian arm and 1.23 on the complex arm. Across the complex arm’s eight onset bins, the median ranged from 1.72 to 1.26 between 2 and 90 diseases per bin, flat apart from the first bin (Fig. 5b). The slope of this ratio on onset was −0.003 (97.5% CI −0.289 to 0.283) on the Mendelian arm and 0.137 (−0.100 to 0.373) on the complex arm (Table 2). The minimum detectable slopes of the realised design were 0.39 and 0.33, respectively, where 0.39 corresponds to the increment falling by a third across the onset range; the design is therefore underpowered for this estimand.

### Balancing selection shows no onset gradient

If late-onset risk alleles were maintained by antagonistic pleiotropy [14, 18], a balancing-selection statistic might carry the onset gradient sought in the purifying-selection scores [41, 51]. Across 5,390 gene-disease rows from 2,911 diseases, the rank correlation between standardised B_2_ [42] and onset was 0.010 (95% CI −0.018 to 0.039), excluding correlations above approximately 0.04. Fitted as a slope across the onset range and expressed in units of the B2 standard deviation of 3.34, it was −0.0008 (95% CI −0.087 to 0.084), excluding gradients above approximately one twelfth of a standard deviation. Median B_2_ was flat across the five onset levels, at 5.65, 5.91, 5.68, 5.40 and 5.90.

### Independent scoring of a genomic language model

Evo 2 was scored on human genes under a stop-codon likelihood rule to provide a model-derived member of the conservation family. It achieved an AUROC of 0.6018 (95% CI 0.587 to 0.617) on the DepMap essentiality [37] dataset across the 16,331 genes with an essentiality label, short of the 0.63 required for inclusion, so it contributes no score to any analysis. The published benchmark reports AUROC 0.66 and area under the precision-recall curve 0.15 for Evo 2 40B on DepMap human gene essentiality, against 0.50 to 0.59 and 0.09 to 0.12 for other genomic language models and 0.65 to 0.71 and 0.13 to 0.21 for four phyloP conservation scores ([9], Extended Data Fig. 3i). Unlike the insertion rule used here, the benchmark scrambles the coding sequence of each gene, preserving dinucleotide composition ([9], Supplementary Information, Section A.3.11), so the deviation from 0.66 reflects differences between two perturbations, applied to different windows and possibly normalised differently. Two observations are independent of that comparison. First, wild-type sequence likelihood alone separated essential genes as well as the likelihood ratio did (paired difference 0.0113, 95% CI −0.0093 to 0.0312), so the perturbation added nothing detectable over baseline sequence typicality. Second, the rule requires a first coding exon long enough to hold the insertion, excluding 3,639 of 20,446 genes. Among the 18,462 genes carrying a DepMap screen, a stricter requirement than holding a label, essential genes were over-represented in the excluded set: 13.75% of 3,186 excluded genes, compared with 9.06% of 15,276 scorable genes (odds ratio 1.60, *P* = 9.2 × 10^−16^). The 16,331 labelled genes and the 15,276 screened scorable genes contain the same 1,384 essential genes. Per-gene scores for 16,794 genes are provided in Additional file 2.

## Discussion

Deep-time conservation and recent human population constraint do not decay at detectably different rates as Mendelian disease onset increases. The point estimate aligns with the direction the analysis plan registered, at 0.024 AUROC units, falling to 0.0035 when somatic and not-yet-assessed associations are dropped, against a design that resolves to 0.049. The interval nonetheless excludes the opposite direction, constraint decaying faster than conservation, beyond 0.011 AUROC units, or 0.033 under that restriction. Three findings qualify this result. First, the gene-class explanation does not account for the contrast. That explanation holds that early-onset disease genes are developmental and pleiotropic by construction, so the relationship between constraint and onset might reflect nothing about the fitness cost of disease. The account requires attenuation of the order of the contrast under conditioning, and attenuation is bounded here at 0.024 against contrasts of 0.23 to 0.50. Two of its three components fail earlier still, since haploinsufficiency and trait-level pleiotropy do not vary with onset among disease genes and so cannot confound an onset interaction. The conditioning half of that evidence bounds confounding of the contrast rather than of either score’s own onset slope, so an account under which gene class steepens both slopes equally is not excluded. Haploinsufficiency also enters as pLI, which is computed from the same loss-of-function counts as LOEUF, so conditioning a constraint score on it is collinear rather than independent. Second, the result depends on the constraint metric, and a result reported on one metric overstates its precision. All three alternative metrics give contrasts excluding zero where *s*_het_ does not, so the conclusion depends on a choice that appears incidental. Paired comparison shows that the estimates are nonetheless inseparable, so the disagreement cannot be settled by argument over which score is preferable. The contrasts under LOEUF and *s*_het_ are close to uncorrelated across disease resamples, at −0.14, although the two scores rank genes alike at 0.82, so the uncertainty comes from which diseases are sampled and is not shared between metrics that agree on genes. Third, the limiting quantity is the number of late-onset Mendelian diseases, 283 genome-wide, which no analytical choice can increase. Constraint metrics are known to differ in their assumptions and in the classes of gene for which they are best powered [1–3, 26, 27], but their consequence for a derived onset contrast has not, to our knowledge, been quantified.

No constraint metric identifies recessive disease genes among controls matched on coding length, expected loss-of-function count and base composition: *s*_het_, LOEUF and missense Z all sit below chance there. This favours the reading on which *s*_het_ behaves as defined rather than underperforming, since a metric blind to these genes would sit at chance rather than below it. Recessive disease accounts for a substantial share of the rare-disease diagnostic yield in paediatric and adult settings [52, 53], and constraint-based filters are routinely applied at the gene level in diagnostic pipelines [4, 5, 7]. The practical implication is that *s*_het_ ranks recessive disease genes below the genome average wherever it is applied as a gene-level prior, so a pipeline using it where recessive disease is in scope needs explicit correction: under a recessive model, a heterozygous constraint prior should not count against a gene, which is a change to how the prior is applied rather than to which metric supplies it. Substituting a loss-of-function depletion measure does not correct this, because the advantage those measures hold against the unmatched pool comes from coding sequence length and expected loss-of-function count, which separate recessive disease genes on their own and from which LOEUF is derived. Against controls matched on them, LOEUF ranks recessive disease genes no higher than *s*_het_ does, so what a pipeline gains by switching metric is the ranking the covariates produce rather than a better reading of constraint. These results are consistent with the allele-level literature. Onset-dependent gradients in risk-allele frequency [17, 18] and in selection signatures below 40 years [19] are measurements on variants; the scores examined here summarise selection across a gene and are dominated by its molecular function. An allele-level gradient does not entail that gene-level scores inherit it at a magnitude detectable across 283 diseases. The theoretical expectation that selection weakens with age [14–16] therefore remains compatible with the absence of a detectable gene-level signature. Two methodological points have wider relevance. Studies adjusting for a gene’s count of associated diseases, therapeutic areas or annotations risk counting the outcome among the covariate’s inputs, a form of multifunctionality bias [47–49] that is invisible when only an annotation-derived measure is available. The problem is compounded by the concentration of biomedical research effort on a minority of genes [50]. Separately, conditioning on annotation-derived covariates requires fitting the comparison on shared rows. Genes lacking biological-process annotation are understudied and rarely disease genes: the dropped rows show a positive rate of 0.024, compared with 0.101 among the kept rows. The deletion is therefore informative about the outcome, although it shifts the fitted sample’s positive rate only from 0.094 to 0.101, since the dropped rows account for a tenth of the total. Part of what then reads as an effect of conditioning is in fact a change of sample.

Several limitations bear on interpretation. The analysis is autosomal, because the gnomAD v4.1 constraint release covers autosomes and the expected loss-of-function count is a matching covariate, removing 211 Mendelian disease genes; the exclusion is not differential with respect to onset, with mean onset position 0.237 retained against 0.213 excluded and Mann-Whitney *P* = 0.81, and costs 6.3% of positive rows, 360 of 5,711. Two controls rest on proxies, tissue specificity measured without conditioning on the affected tissue and the Gene Ontology developmental branch [39] in place of fetal expression; the latter carries measurable dependence on study effort. The conditioning test is informative only for that covariate, since the other two covariates are free of annotation dependence and do not vary with onset, so whether an annotation-independent developmental measure would attenuate more cannot be determined here. Complex-disease onset is defined as the age at first registry record, not age at onset, with a left truncation of 10.2 years for conditions with an English median below 40, which compresses the axis and biases the contrast towards zero. Pleiotropy is measured for adult quantitative traits and will not capture developmental pleiotropy in genes whose severe-variant carriers are absent from the UK Biobank. Finally, Orphanet onset annotation is categorical and curator-assigned [30, 31]; a finer or independently derived onset measure could increase the available resolution, though not the number of late-onset Mendelian diseases.

## Conclusions

Onset stratification of gene-level evolutionary scores is feasible but underpowered at current annotation coverage. The difference between the onset slopes of deep-time conservation and recent human constraint is not resolvable across the 283 late-onset Mendelian diseases available genome-wide, and the standard gene-class explanation for the underlying constraint-onset relationship is not supported by either of its testable components, within the limit that conditioning bounds confounding of the contrast rather than of each score’s slope. Studies that stratify gene-level evolutionary evidence by onset, or by any comparable disease-level axis, should report the minimum detectable effect alongside the estimate [44], since a null on this axis is otherwise uninterpretable, should fit more than one constraint metric, since metrics that rank genes near-identically can yield opposite conclusions about a derived contrast, and should check that no gene-level covariate relies on the same evidence that defines the outcome, as annotation-derived pleiotropy measures routinely do, with an effect that is invisible without an assay-derived comparator. For disease gene prioritisation specifically, *s*_het_ ranks recessive disease genes below the genome average while loss-of-function depletion measures rank them above. The difference is small, precisely estimated and attributable to coding sequence length and expected loss-of-function count: against controls matched on those covariates, every metric ranks recessive disease genes below comparable genes. A pipeline with recessive disease in scope therefore needs an explicit correction rather than a change of metric.

## Supporting information

additional_file_2_evo2_gene_scores

additional_file_3_design_rows.tsv

additional_file_4_disease_table.tsv

additional_file_5_ontology_crosswalk.tsv

additional_file_6_conservation_aggregates.tsv

## List of abbreviations

AUROC: area under the receiver operating characteristic curve
CI: confidence interval
GC: guanine-cytosine
GWAS: genome-wide association study
LOEUF: loss-of-function observed/expected upper bound fraction
pLI: probability of loss-of-function intolerance
*s*_het_: selection coefficient against heterozygotes
TSV: tab-separated values.

## Declarations

### Ethics approval and consent to participate

Not applicable. This study analysed publicly available, de-identified summary-level data and involved no human participants, human material or human data requiring consent.

### Availability of data and materials

All data analysed during this study are publicly available. Source datasets, with releases, retrieval dates and licences, are listed in Additional file 1, Table S1, and the checksum of every retrieved file is recorded in the code repository. Derived tables supporting the conclusions of this article are included as Additional files 2 to 6. Analysis code is available at https://github.com/ahmedanees-m/selection-horizon under the MIT licence and archived on Zenodo (https://doi.org/10.5281/zenodo.23125599). The registered analysis plan is at https://doi.org/10.17605/OSF.IO/CBS8Q [29].

### Authors’ contributions

AAMA conceived the study, designed and implemented the analysis, interpreted the results and drafted the manuscript. EJRN supervised the work, contributed to the interpretation, and critically revised the manuscript for intellectual content. Both authors read and approved the final manuscript.

## Acknowledgements

The authors gratefully acknowledge Vellore Institute of Technology, Vellore, India, for access to computational resources and for the institutional support under which this study was carried out. The authors thank the Orphanet and Orphadata, Open Targets, gnomAD, GENCODE, Gene Ontology, Human Phenotype Ontology, Mondo Disease Ontology, FinnGen and Risteys, GTEx, DepMap and UCSC Genome Browser teams for making their resources publicly available, and Zeng and colleagues [27], Spence and colleagues [40] and Siewert and Voight [42] for releasing their per-gene estimates.

## Additional files

**Additional file 1. Supporting analyses for the onset contrast: design, controls and robustness.** Additional methods and results, with additional figures S1 to S8 and additional tables S1 to S15. Sections cover onset coverage, the crosstab, the complex-disease axis, power, controls, the analysis set, conditioning on gene class, sensitivity to the choice of constraint metric, the genomic language model arm and the balancing-selection arm. (PDF)

**Additional file 2.** Per-gene Evo 2 scores for 16,794 genes. (TSV)

**Additional file 3.** The 55,342 rows of the primary Mendelian design comprise gene, disease, onset position, positive status, and every model term. (TSV)

**Additional file 4.** Disease-level table of onset, inheritance and mean scores for the 2,885 diseases fitted in the primary design. (TSV)

**Additional file 5.** Registry endpoint mappings adjudicated in the Mondo hierarchy, 428 in total, with route, name agreement and verdict. (TSV)

**Additional file 6.** Per-gene conservation aggregates for 21,053 genes, with coding sequence length and GC content. (TSV)

