## additional_file_1 for "Gene-level evolutionary conservation and human constraint decline at comparable rates across the age of onset of Mendelian disease"

### Additional file 1. Supporting analyses for the onset contrast: design, controls and robustness

This file reports the composition of the analysis sets, the power available to detect the contrasts, the control analyses, the conditioning and metric sensitivity analyses, the genomic language model arm and the balancing-selection arm.

Intervals on the contrast, on its components, on the control decays and on the incremental-value slope are Wald intervals from the two-way cluster-robust covariance at alpha 0.025, the error rate the design was powered at, and so have 97.5 per cent coverage. All other intervals are 95 per cent bootstrap percentile intervals.

#### 1. Onset coverage

Age of onset for Mendelian disease is taken from the Orphanet natural history product (v1.3.42, 2026-06-23).

| Quantity | Count |
|---|---|
| ORPHAcodes in the product | 7,790 |
| With any age-of-onset annotation | 6,982 |
| Annotated only as spanning all ages | 868 |
| With an ordinal onset interval | 6,114 |
| With at least one gene association | 4,245 |
| With both an ordinal interval and a gene | 3,524 |

Diseases annotated only as spanning all ages carry no ordinal position and are excluded from the onset axis. They are included at the midpoint of their span in a sensitivity analysis. The gene rows above count any association in the Orphanet gene product. The crosstab in section 2 counts only genes that enter the analysis, which is the narrower set, so its Mendelian onset levels sum to 3,213 rather than 3,524. The seven Orphanet intervals are unevenly occupied, with 521 antenatal and 4 elderly. Antenatal and neonatal are merged because both fall far before first reproduction, and elderly is merged into adult because it holds too few diseases to support a separate level. This gives the five levels used throughout.

#### 2. The crosstab

Diseases and gene-disease pairs by onset level and evidence source.

| Onset | Mendelian diseases | Mendelian pairs | GWAS | ClinVar | Gene burden |
|---|---|---|---|---|---|
| perinatal | 1,749 | 3,263 | 16 | 1,610 | 11 |
| infancy | 477 | 851 | 8 | 399 | 5 |
| childhood | 601 | 1,124 | 23 | 568 | 5 |
| adolescent | 103 | 209 | 5 | 93 | 1 |
| adult | 283 | 484 | 53 | 257 | 17 |
| unassigned | 616 | 1,098 | 9,747 | 6,569 | 1,201 |

The Mendelian arm occupies every level with at least 100 diseases and 200 pairs, so the onset contrast is estimable within it. The complex-disease sources are almost entirely unassigned on this axis, which is expected: Orphanet annotates rare disease and does not cover common disease. They are placed on their own axis in section 3. The adult cell carries the contrast and its composition is therefore reported. Of 283 diseases, 128 fall outside the named categories, 51 are neurodegenerative, 49 are hereditary cancer predisposition, 29 metabolic and 26 cardiac or neuromuscular. Hereditary cancer predisposition is a sixth of the cell, and since its genes are DNA repair genes constrained for reasons unrelated to when the cancer presents, dropping it is a sensitivity analysis, which moves the primary estimate to 0.22.

##### Inheritance composition

| Onset | Dominant | Recessive | Mixed | Other | Dominant share |
|---|---|---|---|---|---|
| perinatal | 488 | 1,043 | 138 | 80 | 0.319 |
| infancy | 111 | 315 | 25 | 26 | 0.261 |
| childhood | 213 | 307 | 46 | 35 | 0.410 |
| adolescent | 50 | 38 | 6 | 9 | 0.568 |
| adult | 155 | 62 | 15 | 51 | 0.714 |

The dominant share more than doubles across the onset range. Since the primary constraint metric is a selection coefficient on heterozygotes, and is close to blind to recessive disease genes, this composition shift would by itself raise the apparent discrimination of constraint at later onset. Mode of inheritance therefore enters the model as a stratifying interaction with each score.

#### 3. The complex-disease axis

Complex-disease onset is the median age at first recorded event in the FinnGen registry (R13), taken from Risteys and binned by decade. 3,666 endpoints carry a usable median after a hundred-case floor. Those medians average 51.1 years, run to a median of 53.6, and occupy every decade. After crosswalking to the Platform disease index, 521 diseases carry an onset decade, and 517 of them have a gene carrying both scores and enter the fitted design.

Two properties of this axis are measured directly.

**Agreement with an independent registry.** Median age at first record for 308 conditions in the English National Health Service was compared with the Finnish values across 161 conditions matched by name. Spearman correlation 0.867, Pearson 0.870. FinnGen records the first event later than the English series in 107 of the 161 conditions, by a median of 2.3 years.

**Left truncation.** A registry that begins recording in 1998 cannot observe the childhood onset of someone already in middle age when it started, so apparent onset is inflated for early-onset conditions. The size of the effect is measurable against the English series: for conditions whose English median is below 40, the Finnish registry records the first event 10.2 years later on average; at 40 and above, 1.3 years. The bias is confined to the part of the axis where the mechanism applies, and it compresses the axis, which biases the onset contrast toward zero.

#### 4. Power

##### The primary contrast

Simulated on the realised design: five onset levels with the disease and pair counts of section 2, ten matched controls per positive, 1,000 replicates per cell, alpha 0.025. The main effect is calibrated from the data, as the pooled marginal discrimination of the constraint score for Orphanet disease genes against matched controls without a score-by-onset interaction. The correlation between the two primary scores, which drives the variance of their difference, is 0.613.

| Stratum | Onset levels | Controls | Minimum detectable effect |
|---|---|---|---|
| All Mendelian | five | matched | 0.035 |
| All Mendelian | five | random | 0.034 |
| All Mendelian | two | matched | 0.041 |
| Dominant only | five | matched | 0.052 |

Units are AUROC-equivalent. These are design-stage figures: the simulation clustered standard errors on disease alone, while the fitted model carries a two-way cluster-robust covariance over disease and gene. On the realised design the simulated standard error of 0.106 logit units understates the fitted 0.147 by a factor of 1.39, of which 1.17 is the second clustering dimension and the remainder follows from the generative assumptions of the simulation. The minimum detectable effects reported in the paper are therefore computed from the realised standard error of each fitted contrast, as that error multiplied by 3.08, the sum of the normal quantiles at alpha 0.025 two-sided and 80 per cent power. For the primary contrast that is 0.45 logit units, or 0.049 AUROC units. The predicted range for the effect if the hypothesis holds is 0.03 to 0.08, so the design resolves only the upper part of that range. An estimate below roughly 0.049 is a bounded null, read against the minimum detectable effect. Two onset levels give a minimum detectable effect of 0.041, because merging levels discards intermediate onset information without adding diseases. Restricting to dominant disease removes the composition confound directly but raises it to 0.052, so inheritance enters as a stratifying interaction instead.

##### The incremental-value slope

Simulated on the realised design, with each disease contributing its own measured baseline discrimination and its own count of positives, and the sampling error of a per-disease AUROC taken from the Hanley-McNeil expression.

| Arm | Diseases above the discrimination floor | Median positives | Minimum detectable slope |
|---|---|---|---|
| Mendelian | 50 | 16 | 0.70 |
| Complex | 278 | 52 | 0.47 |

The slope is on onset scaled to the unit interval, so a simulated slope of 0.70 corresponds to the incremental value halving across the full onset range. These simulated figures are read at zero correlation between the nested AUROCs, which is the conservative corner of the grid. On the same rule as the contrast, the realised standard errors of the fitted slopes give minimum detectable slopes of 0.39 on the Mendelian arm and 0.33 on the complex arm, where 0.39 corresponds to the incremental value falling by a third. The incremental-value slope is not interpreted as a hypothesis test on either figure. The fitted slopes are −0.003 with an interval of −0.289 to +0.283 on the Mendelian arm, and +0.137 with an interval of −0.100 to +0.373 on the complex arm. No decay was detected, and what could have been detected is a decay of 0.39 and 0.33 respectively. As a descriptive quantity, the ratio has a median of 1.17 on the Mendelian arm and 1.23 on the complex arm, so the evolutionary scores add roughly a fifth to the lift of a model built without them. Across the complex arm's eight onset bins the median runs 1.72, 1.32, 1.16, 1.24, 1.21, 1.26, 1.20 and 1.26, on between 2 and 90 diseases each, which is flat apart from the first bin's two diseases. The Mendelian arm's later bins hold three or four diseases each and support no statement about shape. Two assumptions bear on these figures. The sampling errors of the two nested models are treated as independent, which they are not, since one model contains the other and both are fitted on the same rows; this overstates the noise, and under a correlation of 0.5 the two values are 0.48 and 0.34. The baseline level of the ratio must be set because the variance of its logarithm depends on it, and it cannot be estimated without fitting the model the estimand comes from; the table uses 1.30, and the full grid is given in Table S7.

#### 5. Controls

Four control analyses, all fitted on the Mendelian arm. Each decay control is fitted on its own score with both evolutionary scores removed from the design, so that no control is conditioned on the quantities it is meant to be independent of.

| Control | Type | Decay | 97.5% interval |
|---|---|---|---|
| Coding sequence length | Negative | -0.019 | -0.118 to 0.080 |
| Tissue specificity | Negative | 0.032 | -0.168 to 0.232 |
| Developmental annotation | Positive | 0.154 | 0.043 to 0.264 |
| Onset permuted within therapeutic area | Placebo | -0.009 (mean) | -0.278 to 0.253 (central 95% of draws) |

Neither negative control carries an onset gradient. The positive control decays with an interval excluding zero. Over 8,000 draws the permutation null is centred at −0.009, which is 0.06 of its own standard deviation from zero, with a Monte Carlo standard error on that centre of 0.011 standard deviations. The null is therefore centred on zero to within a tenth of its spread. The width of the null is informative independently of its centre: its central 95 per cent span implies a standard deviation of 0.136 logit units, against the 0.147 standard error of the fitted contrast. The permutation and the sandwich agree on the precision of the design by independent routes. Tissue specificity is measured across all tissues, without conditioning on the tissue a disease affects, because none of the sources provides a disease-to-tissue mapping. Developmental status is the share of a gene's biological-process terms in the developmental branch of the Gene Ontology; fetal expression, the direct measure, is available as alignments rather than as a per-gene matrix. The positive control demonstrates that the onset axis resolves gene class. Because developmental genes are also pleiotropic and haploinsufficient, it does not distinguish gene class from fitness cost; that separation is addressed in Section 7. Coding sequence length is also one of the three covariates positives and controls are matched on, so that control is bounded by construction and tests only the variation left within a decile.

#### 6. Analysis set

The constraint metrics released with gnomAD v4.1 cover the autosomes and carry no genes on chromosome X or Y; four canonical X-linked disease genes, DMD, F8, FMR1 and AR, are absent from the release. The v2.1.1 release carries 768 X and 25 Y genes. Expected loss-of-function count is one of the three covariates positives are matched on, and it derives from the v4.1 release, so genes on the sex chromosomes carry no value for it and leave the design. 211 Mendelian disease genes that otherwise qualify are excluded on this basis. The analysis is therefore autosomal, and X-linked disease is absent from the inheritance stratification as well. X-linked disease is enriched for early-onset developmental conditions, so a loss that differed by onset would flatten the contrast at the early end. The loss does not differ by onset. The table counts the positives whose gene carries both primary scores, which are the rows that would otherwise enter the design.

| Onset | Retained | Excluded | Excluded share |
|---|---|---|---|
| perinatal | 2,987 | 189 | 0.059 |

| Onset | Retained | Excluded | Excluded share |
|---|---|---|---|
| infancy | 732 | 80 | 0.099 |
| childhood | 1,003 | 63 | 0.059 |
| adolescent | 180 | 11 | 0.058 |
| adult | 449 | 17 | 0.036 |

The retained column sums to 5,351, which is 29 more than the 5,322 positives the design fits: 13 are disease-gene pairs that Orphanet lists twice and the design counts once, and 16 are genes with no expected loss-of-function count, which is a matching covariate. Mean onset position 0.237 among retained positives against 0.213 among excluded ones, Mann-Whitney p = 0.81. The loss is 360 of 5,711 positive rows, 6.3 per cent, and is spread across the axis, so it costs precision without tilting the contrast.

#### 7. Conditioning on gene class

The gene-class account holds that early-onset disease genes appear constrained because they are developmental, pleiotropic and haploinsufficient, so conditioning on gene class should attenuate the contrast toward zero. It fails on all three components, by two kinds of evidence. The conditioning half of that evidence bounds confounding of the contrast rather than of either score's onset slope, so an account under which gene class steepens both slopes equally is not excluded. The developmental component is the one this design can test by conditioning, and conditioning does not attenuate, with the attenuation left open bounded at 0.024 against contrasts of 0.229 to 0.495. The other two fail at their premise: neither pleiotropy nor haploinsufficiency varies with onset among disease genes, so neither can be producing a gradient in the first place.

##### Row sets

Five row sets are in use. The unconditioned contrast differs across them, so every figure below names the set it was fitted on.

| Row set | What it holds | Rows | Contrast |
|---|---|---|---|
| Primary, Mendelian arm | Every gene carrying s_het and the conservation metric | 55,342 | 0.2275 |
| Mediation, shared rows | The rows every mediation specification can be fitted on, so conditioning is not read together with a change of coverage | 50,138 | 0.2471 |
| Common gene set, all rows | Only genes carrying all three constraint metrics, so that swapping the metric is the only thing that varies | 57,090 | 0.2017 |
| Common gene set, retained covariates | As above, restricted to the rows the three retained covariates cover | 51,616 | 0.2273 |
| Common gene set, all four covariates | As above, including disease breadth, so the component split compares like with like | 51,585 | 0.2295 |

The five contrasts span 0.2017 to 0.2471, a spread of 0.045, against a sampling interval on the primary estimate of roughly 0.33 either way. The restrictions differ from one another by about a seventh of the uncertainty on any one of them, so the choice of row set does not drive the estimate. The conditioning movement differs between the two designs for the same reason: +0.026 on the mediation rows, from 0.247 to 0.273, and +0.006 on the common gene set. The second carries an interval of −0.006 to +0.049 which contains the first, so the two are consistent and the difference is the gene set rather than the specification.

##### Sample composition across specifications

The covariates do not cover every gene, so a specification entering them is fitted on fewer rows than one that does not. A movement measured across that gap combines a change of sample with the effect of conditioning. For the four-covariate block, where the rows fall from 57,090 to 51,585, the two parts separate as follows.

| Constraint metric | Contrast, all rows | Contrast, shared rows | Change of sample | Change from conditioning | Total movement |
|---|---|---|---|---|---|
| s_het | 0.202 | 0.229 | +0.028 | +0.046 | +0.074 |
| Missense Z | 0.375 | 0.384 | +0.009 | +0.034 | +0.043 |
| LOEUF | 0.481 | 0.495 | +0.015 | +0.048 | +0.063 |

On the primary metric the sample accounts for 38 per cent of the total. The deletion is not ignorable. Of the 5,505 rows dropped, 5,166 go to the developmental covariate on its own, which is a share of biological-process terms and undefined for a gene that has none. A gene with no process annotation is an understudied gene, and understudied genes are rarely disease genes: the dropped rows carry a median of zero process terms against eight among the rows kept, and a positive rate of 0.024 against 0.101. The deletion is therefore informative about the outcome. Its effect on the fitted sample is smaller than that ratio suggests, because the dropped rows are a tenth of the total: the positive rate rises from 0.094 across all 57,090 rows to 0.101 across the 51,585 kept. Every conditioning estimate in this section is fitted on the rows its specifications share, with the contrast on all rows reported beside it.

##### Covariate-wise conditioning

Each covariate was entered on its own and the block jointly, on identical rows, bootstrapped over 400 resamples of diseases.

| Covariate | Source | s_het | LOEUF | Missense Z |
|---|---|---|---|---|
| Haploinsufficiency | gnomAD allele counts | +0.000 [-0.010, +0.022] | +0.035 [+0.021, +0.055] | +0.024 [+0.013, +0.045] |
| Developmental share | Gene Ontology curation | +0.007 [-0.007, +0.038] | -0.013 [-0.020, +0.004] | +0.002 [-0.024, +0.015] |
| Trait breadth | UK Biobank effect estimates | -0.001 [-0.005, +0.005] | -0.002 [-0.004, +0.001] | -0.002 [-0.005, +0.005] |
| Disease breadth | Open Targets evidence sets | +0.054 [+0.019, +0.087] | +0.032 [-0.004, +0.044] | +0.019 [-0.013, +0.046] |
| The three retained | | +0.006 [-0.006, +0.049] | +0.020 [+0.009, +0.050] | +0.025 [-0.002, +0.052] |
| All four together | | +0.046 [+0.020, +0.101] | +0.048 [+0.011, +0.078] | +0.034 [-0.001, +0.081] |

Disease breadth is not used as an adjustment covariate. It is assembled from the Open Targets evidence sets, one of which is the Orphanet set that supplies the Mendelian positives, so the covariate counts among a gene's therapeutic areas the association that makes it a positive here, and its construction includes the outcome. On the primary metric it moves the contrast by 0.054, against at most 0.007 for any other covariate, which is what a covariate carrying the outcome would do. It is reported on its own above.

##### Attenuation bounds

The three retained covariates were each entered separately and none attenuates the contrast. Across those nine cells and the three joint fits, the lowest interval bound anywhere is −0.024, so an attenuation larger than about 0.024 is excluded in every one of them. The contrasts being conditioned run from 0.229 to 0.495. The account requires gene class to produce the contrast, which requires an attenuation of the order of the contrast itself, and what the data leave open is a tenth of that. The bound rests on the three covariates being free of disease annotation and each entered separately. Haploinsufficiency is pLI from gnomAD allele counts, with no disease annotation involved. Trait breadth is the effective number of traits from the Spence per-gene effect estimates across UK Biobank quantitative traits. Developmental share is Gene Ontology curation, which is discussed below. Freedom from disease annotation is not sufficient on its own. The next section asks whether each covariate varies with onset, since one that does not could not have attenuated the contrast and its null movement would carry no information. Only the developmental share does, and the −0.024 bound is exactly its three cells. On the mediation specification, fitted on the 50,138 rows every specification shares, the contrast is 0.247 unconditioned and 0.273 conditioned on gene class and trait breadth.

**Covariate-onset gradients**

A covariate can confound the score-by-onset interaction only if it varies with onset. One that is flat along the axis could not have attenuated the contrast whatever its relation to the scores, so a null movement from it is uninformative. Rank correlation with onset, resampled over diseases:

| Covariate | Among disease genes | Across all design rows |
|---|---|---|
| Developmental share | -0.083 [-0.121, -0.048] | -0.020 [-0.031, -0.009] |
| Haploinsufficiency | +0.014 [-0.034, +0.062] | +0.006 [-0.006, +0.017] |
| Trait breadth | +0.011 [-0.015, +0.037] | +0.003 [-0.006, +0.012] |
| Disease breadth | -0.011 [-0.041, +0.023] | +0.006 [-0.011, +0.028] |

The design carries ten matched controls for every positive, drawn without regard to these covariates, so a gradient among disease genes is diluted about elevenfold across the whole design. The counter-hypothesis is a claim about disease genes, so the first column is the one it is made on. Only the developmental share varies with onset. The counter-hypothesis therefore separates into one component that conditioning can test and two that fail at their premise.

**The developmental half is tested.** The developmental share falls with onset among disease genes on an interval clear of zero, in the direction the account predicts, so it is a covariate that could have carried the contrast. Conditioning on it does not attenuate: +0.007, −0.013 and +0.002 across the three metrics, every interval containing zero, and the −0.024 bound above is these three cells.

**The pleiotropy half fails at its premise, for the pleiotropy that was measured.** The account asserts that early-onset disease genes are more pleiotropic. Trait breadth, the pleiotropy measure not built from disease annotation, does not vary with onset among disease genes at all: +0.011, bounded within 0.037 either way. So the assertion is refuted directly, and the conditioning result on that covariate is an uninformative test, not support. Trait breadth is the effective number of traits from the Spence per-gene effects across UK Biobank, which measures adult quantitative-trait pleiotropy: carriers of severe early-onset variants are largely absent from that cohort, and the traits measured are not developmental ones. The account's sense of pleiotropy is nearer to developmental or multi-organ action. So what is refuted at the premise is the quantitative-trait version of the claim. The developmental version of the same idea is carried by the developmental share, which does vary with onset and which fails the conditioning test instead.

**The haploinsufficiency half fails the same way.** pLI is flat along onset among disease genes at +0.014, bounded within 0.062. The crosstab runs against the account here as well, since the dominant share rises from 0.319 to 0.714 with later onset rather than falling. The account therefore fails on all three of its components, by two kinds of evidence rather than one, and the conditioning test carries only the component for which it is informative.

**Interpretation of the conditioning movements**

Conditioning on a common cause of onset and constraint removes confounding. Conditioning on a variable that is partly a descendant of disease-gene status can create association instead. With disease breadth excluded, two of the nine remaining cells have an interval clear of zero, and both are haploinsufficiency: +0.035 [+0.021, +0.055] on LOEUF and +0.024 [+0.013, +0.045] on missense Z, with the joint fit that carries the first at +0.020 [+0.009, +0.050]. Both have an alternative reading. pLI and LOEUF are computed from the same gnomAD loss-of-function counts, observed against expected, so conditioning LOEUF on pLI is conditioning a constraint score on a near-duplicate of itself. A movement there is as readily collinearity as gene-class confounding, and the causal graph does not distinguish the two because it classifies haploinsufficiency as a gene-class marker rather than as a constraint measure. Missense Z is the missense counterpart in the same dataset, which is a weaker form of the same objection but the same objection. There is a further reason not to read either as gene class being removed. pLI does not vary with onset among disease genes, as the section above reports, so it is not a covariate that could have been confounding an onset interaction in the first place. The counter-hypothesis fails, and no reading is available in which gene class suppresses the relationship. The developmental covariate moves no estimate. Every interval contains zero and no point estimate exceeds 0.013 in size. It correlates 0.296 with a gene's biological-process term count across 16,832 genes, against 0.369 for the raw developmental count, so taking the share reduced the dependence on study effort without removing it. The concern that Gene Ontology annotation is partly downstream of disease-gene status is therefore real and immaterial here, because the covariate it applies to does not move the estimate either way.

**Scope and limitations**

The conditioning test is informative for one covariate only, the developmental share, because it is the only one that varies with onset among disease genes. That covariate is also the one with measurable dependence on study effort, correlating 0.296 with a gene's process-term count. The two covariates free of that dependence, pLI and trait breadth, do not vary with onset, so they cannot test the hypothesis by conditioning at all. The 0.024 bound therefore rests on a covariate that is partly a record of what has been studied. Whether a developmental measure free of annotation effects would attenuate more cannot be determined here; that requires a fetal transcriptome, which is not available as a per-gene table. Section 5 describes the developmental measure. The premise-level part of the refutation is not affected. It rests on the two covariates that are free of the dependence, and on the finding that they are flat along the onset axis among disease genes.

**Relation to multifunctionality bias**

Disease breadth behaves as multifunctionality bias predicts. Gillis and Pavlidis showed that multifunctionality rather than association is a primary driver of gene function prediction, that the degree of multifunctionality alone predicts function strongly, and that the bias has to be controlled for if predictions are to be interpreted. A later study of autism gene prioritisation scores found them correlating with physical node degree and with publication count at Spearman 0.34, and stated the mechanism plainly: genes become studied as they are associated with disease, and they then accumulate functional and physical annotations. The 0.296 measured here between the developmental share and process term count sits in that range. That study left one question open, whether top-ranked genes really are more multifunctional or only appear to be through biases in research and in patterns of annotation. The two pleiotropy covariates here bear on this question for this construct. Both were entered to measure one thing, on the same genes in the same model. Trait breadth, built from assay data, moves the contrast by less than 0.002 under every metric, on the tightest intervals in the table. Disease breadth, built from disease annotation, carries the whole of the movement on the primary metric. Two measures of pleiotropy disagreeing that completely is consistent with the second reflecting annotation bias rather than pleiotropy. Trait breadth and disease breadth are not two measures of one construct, however, and a covariate drawing on the evidence that defines the outcome would move the estimate whatever pleiotropy did. That comparison is not available in a design whose only measure of pleiotropy is the annotation-derived one, which is the usual case.

#### 8. Sensitivity to the choice of constraint metric

Whether the contrast excludes zero depends on which constraint metric is used. It does not follow that the metrics measure different things, and the stronger reading is not supported by this design. Each metric is fitted as its own arm, so the estimate and the interval below come from the same design in every row. The common gene set used later in this section gives slightly different point estimates and is used only where the three have to be compared with each other.

| Constraint metric | Contrast | 97.5% interval | Rows |
|---|---|---|---|
| s_het, primary metric | 0.227 | -0.102 to 0.557 | 55,342 |
| Missense Z | 0.377 | +0.133 to 0.621 | 56,942 |
| LOEUF | 0.466 | +0.280 to 0.652 | 56,942 |
| LOEUF, gnomAD v2.1.1 | 0.475 | +0.308 to 0.643 | 55,442 |

Every alternative excludes zero and the primary metric does not, so the conclusion depends on the metric.

##### Paired comparison between metrics

The estimates are fitted on the same genes, the same positives and the same design, so they are compared as a difference over the same disease resamples rather than through their marginal intervals.

Bootstrapped over 400 resamples of diseases, on the common gene set:

| Comparison | Observed difference | 95% interval |
|---|---|---|
| LOEUF minus s_het | +0.279 | -0.037 to +0.551 |
| Missense Z minus s_het | +0.174 | -0.017 to +0.391 |

Neither excludes zero, though both come close. So the metrics are not shown to differ. The choice between them decides whether the contrast excludes zero, while the data cannot say that the underlying quantities are different. The result is about power, not about the scores. An onset-stratified constraint contrast at this sample size cannot separate constraint metrics from one another, so a result reported on one metric carries a sensitivity that its own interval does not show. The resamples say what would have to change. Across the 400 disease resamples the contrast under LOEUF correlates −0.14 with the contrast under s_het, and the contrast under missense Z correlates 0.59 with it, against rank correlations of 0.82 and 0.63 between the scores themselves. The bootstrap standard deviation of the contrast is 0.130 under s_het, 0.070 under LOEUF and 0.085 under missense Z. The sampling error of each contrast therefore comes from which diseases fall into the resample and is largely specific to the metric, even between metrics that rank genes alike. The limiting quantity is the number of late-onset diseases, which is 283 in the Mendelian arm, and no change of constraint metric addresses that.

**Decomposition of the contrast**

The rest of this section concerns the pattern in the point estimates. Everything in it is conditional on that pattern being real, which the paired comparison above does not establish. The contrast is a difference of two decays, and conservation is held at the same metric in all three rows, so the components locate the pattern.

| Constraint metric | Conservation decay | Constraint decay | Contrast |
|---|---|---|---|
| s_het | 0.346 | +0.144 | 0.202 |
| Missense Z | 0.414 | +0.039 | 0.375 |
| LOEUF | 0.403 | -0.078 | 0.481 |

The constraint components have opposite signs at the point estimate: s_het falls with onset, missense Z is flat, LOEUF rises. Those components inherit the same imprecision as the contrasts they make up and are not separable either. The conservation component moves across a range of 0.068. This is expected, since the conservation coefficient is fitted jointly with the constraint score and the three metrics share variance with it to different degrees.

The scores themselves agree, and the agreement does not vary along the axis:

| Stratum | s_het and LOEUF | s_het and missense Z | LOEUF and missense Z |
|---|---|---|---|
| All genes | -0.821 | 0.626 | -0.733 |
| perinatal | -0.807 | 0.596 | -0.722 |
| infancy | -0.878 | 0.717 | -0.749 |
| childhood | -0.820 | 0.656 | -0.738 |
| adolescent | -0.755 | 0.600 | -0.753 |
| adult | -0.816 | 0.645 | -0.743 |

Spearman correlations, with LOEUF inverted by construction so that a negative value is agreement. Three accounts of the point-estimate pattern were tested, and none explains it. It is not differential coverage, since the common gene set does not close it. It is not gene-class content in the estimator's prior: on the shared row set, conditioning on gene class moves s_het by +0.007 against +0.027 for the largest direct measure, so it moves least where the account needs it to move most. It is not inheritance composition, since LOEUF's dominant-to-recessive gap is narrower where that account needs it wider. Every movement under conditioning is between 0.006 and 0.027, against a point-estimate spread of 0.28 that is itself not distinguishable from zero. So, conditioning of the magnitude available in this adjustment set could not account for the pattern even if the pattern were established. The pattern in the point estimates therefore has no demonstrated explanation, and is not itself established.

**Inheritance composition**

Mode of inheritance shifts along the onset axis, with the dominant share rising from 0.319 to 0.714 across the five levels, and the constraint metrics discriminate dominant disease genes better than recessive ones. If a metric with no feature prior were less able to identify recessive genes, its discrimination would rise across an axis that becomes more dominant, and that would produce the observed pattern. The prediction is that LOEUF, a heterozygous depletion measure with no prior, should show a wider gap between dominant and recessive genes than s_het does. It does not.

| Constraint metric | Dominant only | Recessive only | Gap |
|---|---|---|---|
| s_het | 0.723 | 0.487 | +0.236 |
| LOEUF | 0.725 | 0.533 | +0.192 |
| Missense Z | 0.682 | 0.521 | +0.160 |

Discrimination of Mendelian disease genes within each stratum, on the genes carrying all three metrics. The account needs LOEUF's gap to be wider than s_het's and it is narrower, so it fails on its point prediction. Against matched controls the same ordering holds, with gaps of 0.217 for s_het, 0.199 for LOEUF and 0.162 for missense Z.

##### Discrimination by inheritance stratum

The stratified figures carry a separate result that does not depend on the contrast at all. They are computed against two reference sets, the pool of scored genes and controls matched on the three covariates of the primary design, because the pool carries those covariates inside it.

| Stratum | Metric | Discrimination | 95% interval |
|---|---|---|---|
| Recessive | s_het | 0.487 | 0.476 to 0.498 |
| Recessive | LOEUF | 0.533 | 0.521 to 0.544 |
| Recessive | Missense Z | 0.521 | 0.509 to 0.534 |
| Dominant | s_het | 0.723 | 0.707 to 0.737 |
| Dominant | LOEUF | 0.725 | 0.710 to 0.740 |
| Dominant | Missense Z | 0.682 | 0.665 to 0.698 |

Bootstrap intervals over 1,000 resamples, 2,173 recessive and 1,218 dominant disease genes against a pool of 17,434 scored genes. The between-metric difference is the well determined part. Compared paired on the same genes, LOEUF exceeds s_het on recessive disease genes by 0.046, with an interval from 0.037 to 0.055. s_het's own distance from chance is 0.013, which is resolved only because the sample is large and is negligible in use. Two readings of that divergence were available before the comparison was repeated against matched controls. On the first, s_het is behaving as defined. It is a selection coefficient on heterozygotes, recessive disease genes have unaffected heterozygotes, and an estimator that ranks them below average is reporting something true about heterozygous selection. The number needing explanation is then LOEUF at 0.533: why a heterozygous loss-of-function depletion measure finds recessive disease genes more depleted than average. On the second, the feature prior is pushing recessive genes the wrong way, since they are enriched for restricted-expression enzymes whose feature profile reads as unconstrained.

The matched comparison answers the question the first reading raises. The pool carries the matching covariates inside it. Recessive disease genes have a median coding sequence of 1,662 bases against 1,359 for the pool and a median expected loss-of-function count of 50.2 against 35.4, and LOEUF is a function of that count. Against ten controls per gene drawn from genes causing no Mendelian disease in the same stratum of coding sequence length decile, expected loss-of-function count decile and GC decile, LOEUF falls to 0.450 and s_het to 0.457. The LOEUF advantage does not survive and reverses, to −0.0067 with an interval from −0.0136 to −0.0004. Missense Z stays marginally above s_het, at +0.0150 from 0.0049 to 0.0246. All three metrics place recessive disease genes below comparable genes. On dominant disease genes the matched comparison separates the metrics in the other direction, with s_het at 0.674 above LOEUF at 0.650 and missense Z at 0.634, which is what a selection coefficient on heterozygotes should do.

| Stratum | Metric | Scored pool | Matched controls |
|---|---|---|---|
| Recessive | s_het | 0.487 | 0.457 |
| Recessive | LOEUF | 0.533 | 0.450 |
| Recessive | Missense Z | 0.521 | 0.472 |
| Dominant | s_het | 0.723 | 0.674 |
| Dominant | LOEUF | 0.725 | 0.650 |
| Dominant | Missense Z | 0.682 | 0.634 |

Discrimination against each reference set. Matched controls are ten per gene, from genes causing no Mendelian disease in the same stratum of coding sequence length decile, expected loss-of-function count decile and GC decile.

The operational consequence is the same under either reading and does not depend on resolving it. A gene prioritisation that uses s_het where recessive disease is in scope will rank those genes below average genes. Swapping in a direct depletion measure changes that ranking through coding length and expected loss-of-function count rather than through constraint, so it is not a correction.

The matching holds all three covariates close. Standardised mean differences after matching are 0.074 on coding sequence length, 0.097 on expected loss-of-function count and 0.006 on GC content for recessive genes, against 0.309, 0.404 and −0.042 for the same genes against the unmatched pool. Matching supplied 9.62 controls per recessive gene and 9.65 per dominant gene. 154 recessive and 76 dominant genes sit in strata holding fewer than ten eligible controls and take what their stratum has. The direction holds under five controls per gene, under twentieth-width strata in place of deciles, and under a within-set estimator that scores each gene only against its own controls, which give LOEUF minus s_het between −0.007 and −0.022 on recessive genes and between −0.005 and −0.038 on dominant ones.

The arm that speaks to the same question is the three-way interaction, which lets the onset slope differ by mode rather than assuming it common. Both columns are fitted on the genes carrying all three metrics, so the comparison is between specifications and not between samples.

| Constraint metric | Two-way | 97.5% interval | Three-way | 97.5% interval |
|---|---|---|---|---|
| s_het | 0.202 | -0.136 to 0.539 | 0.465 | -0.207 to 1.136 |
| Missense Z | 0.375 | +0.134 to 0.617 | 0.479 | +0.056 to 0.903 |
| LOEUF | 0.481 | +0.297 to 0.664 | 0.401 | -0.093 to 0.895 |

The point estimates converge, with the spread across metrics falling from 0.279 to 0.079, but the intervals do not support reading anything into that. They run 0.85 to 1.34 wide against 0.68 or less for the two-way estimates on the same rows. On the primary arm the same comparison is explicit: the two-way estimate is 0.227 with an interval of −0.102 to 0.557, and the three-way is 0.429 with an interval of −0.220 to 1.077. The three-way interval contains the two-way estimate, so the data are consistent with no sensitivity to the mode specification at all. The point estimate moves with the specification, and this design cannot determine whether that movement is real.

#### 9. The genomic language model arm

Evo 2 was scored on human genes under a stop-codon likelihood rule specified for this analysis. Inclusion in the conservation family required an AUROC for DepMap essential genes within 0.03 of the published 0.66. The measured 0.6018 falls short of that, so Evo 2 contributes no score to any analysis. The measurements bear on how genomic language models are evaluated and are reported for that reason.

##### Scoring procedure

The published evaluation of Evo 2 includes a human essentiality benchmark, in which the model's response to a scrambled coding sequence separates genes that are essential in cell lines from those that are not. The 40B model reaches an AUROC of 0.66 there, with an area under the precision-recall curve of 0.15, against 0.50 to 0.59 and 0.09 to 0.12 for other genomic language models and 0.65 to 0.71 and 0.13 to 0.21 for four phyloP conservation scores (Extended Data Fig. 3i of the published evaluation). The benchmark scores the difference in log likelihood between the coding sequence of a gene and a scramble of it that preserves dinucleotide composition, with no flanking sequence and only the first 8,192 bases of longer genes (Supplementary Information of the published evaluation, Section A.3.11). The rule below was specified for this analysis and differs from the benchmark in its perturbation, which substitutes a stop codon near the start of the coding sequence where the benchmark scrambles the whole sequence. It also scores an 8 kb genomic window centred on the coding start where the benchmark scores the coding sequence alone, and its normalisation may differ. Any of the three could account for a difference of the size seen, so the comparison with 0.66 is a comparison between two perturbations. The hosted 40B model returns vocabulary logits at every position of a supplied sequence, which is what a likelihood ratio needs. For each gene with a canonical coding start, an 8 kb genomic window centred on that start was scored, then scored again with a stop codon replacing the codon at nucleotide offsets 15, 30 and 45. The score is the mean difference in sequence log likelihood across the three positions, normalised by window length. Of 16,807 genes submitted, 16,794 returned a score, and 16,331 of those carry a DepMap essentiality label. The returned likelihoods separate real from scrambled sequence: the ACTB coding sequence scores −0.276 in log likelihood per base, a composition-matched shuffle of it −1.377, and uniform random sequence of the same length −1.411.

##### Results

**Discrimination is below the published figure.**

| Quantity | Value |
|---|---|
| AUROC across the 16,331 scored genes with a label | 0.6018 |
| Bootstrap 95 per cent interval | 0.587 to 0.617 |
| Published figure | 0.66 |
| Coding length alone | 0.499 |
| GC content alone | 0.419 |
| Both baselines together | 0.580 |

The interval does not contain the published value. It lies above chance and above what the simple sequence properties reach.

**The perturbation adds nothing detectable.** The wild-type sequence log likelihood, taken on its own with no stop codon involved, separates the same genes at 0.6132 against 0.6018 for the stop-codon ratio. Compared paired on the same 16,331 genes, the difference is +0.0113 with a bootstrap interval from −0.0093 to +0.0312. The interval contains zero, so the wild-type likelihood is not established as the better of the two. The likelihood ratio does not improve on the wild-type likelihood, however, and the perturbation the rule is built on contributes nothing detectable at this sample size. The score therefore reflects how typical the gene's genomic neighbourhood looks to the model rather than the model's response to a loss-of-function edit. Typicality correlates with sequence conservation, which correlates with essentiality, so an apparently functional prediction reduces to a conservation-like quantity.

**The summation window does not dilute the signal.** The edit changes three bases inside 8,192. Restricting the likelihood difference to the 200 bases beginning at the edit, on the 292 of 300 rescored genes carrying a DepMap label, moves discrimination by −0.0016 with a 95 per cent interval from −0.029 to +0.024, and the two scorings rank genes at a Spearman correlation of 0.959. Per-position values show why. Across 200 genes and three stop positions, 88.0 per cent of the signed likelihood difference sits in the first 200 bases after the edit, 8.6 per cent in the next 800, and 3.4 per cent in the remaining seven kilobases. Positions before the edit contribute a signed total of −10.5 against −40,415 overall, as an autoregressive model requires.

**The gene sets are not comparable.** Placing a stop codon at a fixed early offset requires a first coding exon long enough to hold it, which excludes 3,639 of 20,446 genes. Those genes are not a random sample with respect to the label being scored.

| Gene set | DepMap screened | Essential | Share |
|---|---|---|---|
| Scorable | 15,276 | 1,384 | 0.0906 |
| Excluded | 3,186 | 438 | 0.1375 |

Essential genes are over-represented among the excluded, odds ratio 1.60 against the scorable set, $p = 9.2e\text{-}16$, so the scored genes differ in composition from the full screened set.

##### Gene set composition

The excluded genes cannot be scored, so their contribution cannot be measured. Its direction can be bounded by asking whether discrimination depends on first coding exon length among the genes that can be scored.

| First coding exon | Genes | Essential | AUROC |
|---|---|---|---|
| 48 to 88 bases | 4,161 | 406 | 0.6513 |
| 89 to 154 | 4,016 | 477 | 0.5467 |
| 155 to 364 | 4,071 | 351 | 0.5624 |
| 365 and above | 4,083 | 150 | 0.5438 |

Discrimination is highest in the quartile with the shortest first coding exons and roughly flat across the rest. The excluded genes sit below that quartile. If the pattern continues past the observed range, including them would raise the AUROC rather than lower it, so 0.6018 would understate what the full gene set gives and the gap to the published figure would be overstated. This is an extrapolation beyond the observable range, and it bounds a direction, not a magnitude. A prevalence-matched resample of the scorable genes would not change the figure. AUROC is a rank statistic over positive and negative pairs and is invariant to class prevalence, so matching the essential share of the full screened set returns the same number by construction. What is unknown is the excluded genes' scores, not their prevalence.

##### Limitations
The claim is narrow. Under a stop-codon likelihood rule specified here, at the model size of the published figure, human gene essentiality separates at about 0.60, and the perturbation that rule is built on adds nothing over the model's baseline surprise at the sequence. The second part does not depend on the published protocol: within this rule, the likelihood ratio does not improve on the wild-type likelihood alone. The measurement does not bear on whether the published result is correct. Three things separate it from the published one: the perturbation, since the benchmark scrambles the coding sequence where this rule substitutes a stop codon; the sequence scored, an 8 kb genomic window against the coding sequence alone; and the gene sets, which differ and cannot be reconciled without the published per-gene scores. The effect of the exclusion is bounded in direction rather than measured. Evo 2 is one candidate member of the conservation score family, and the analysis proceeds on three alignment-based tracks. Nothing in this section enters the primary contrast or the mediation.

##### Code and data
The scoring code, its evaluation against DepMap and the writer for the per-gene table are part of the analysis repository. The scores, the per-position likelihood arrays and the evaluation record are regenerable from the pinned sources. The per-gene scores are provided as Additional file 2, whose header records the model, the window, the insertion rule, the exclusion and its enrichment, and the discrimination figure.

#### 10. The balancing-selection arm
Standardised B2 aggregated per gene across three populations, with the HLA region and the olfactory receptor family excluded throughout. The question is whether a balancing-selection statistic shows the onset gradient that the purifying statistics were tested for. It does not, and the size of the absence is bounded. Across 5,390 gene-disease rows on the Mendelian arm, the rank correlation between B2 and onset is +0.010 with a bootstrap interval of −0.018 to +0.039, so a rank correlation larger than about 0.04 in either direction is excluded. Fitted as a slope across the full onset range and expressed in units of the B2 standard deviation of 3.34, it is −0.0008 with an interval of −0.087 to +0.084. A gradient larger than about a twelfth of a standard deviation across the whole axis is therefore excluded. Median B2 by onset level, for reference: 5.65 at perinatal, 5.91 at infancy, 5.68 at childhood, 5.40 at adolescent and 5.90 at adult. The arm uses standardised B2 alone, since Tajima's D and allele-age discordance have no per-gene release.

#### Additional figures

**Figure S1** Power for the primary contrast

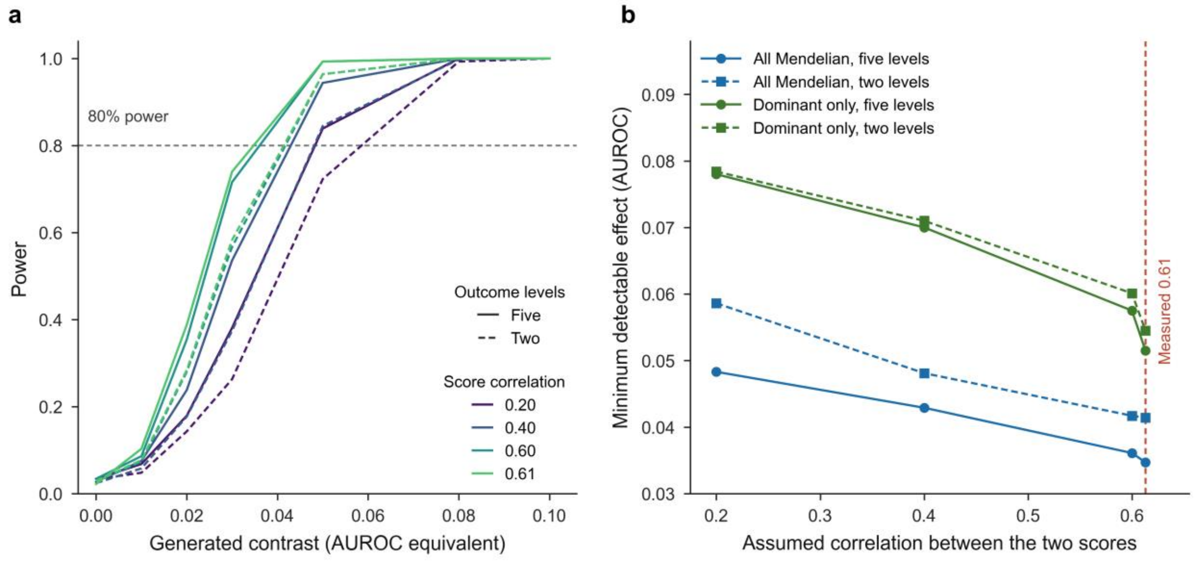


**a** Power against the generated contrast, in AUROC-equivalent units, for each assumed correlation between the two scores and for the five-level and two-level outcomes. The dashed line marks 80% power. **b** Minimum detectable effect against the assumed correlation, for all Mendelian diseases and for dominant disease only, with the correlation measured in the data marked at 0.61.

**Figure S2** Power for the incremental-value slope on both arms

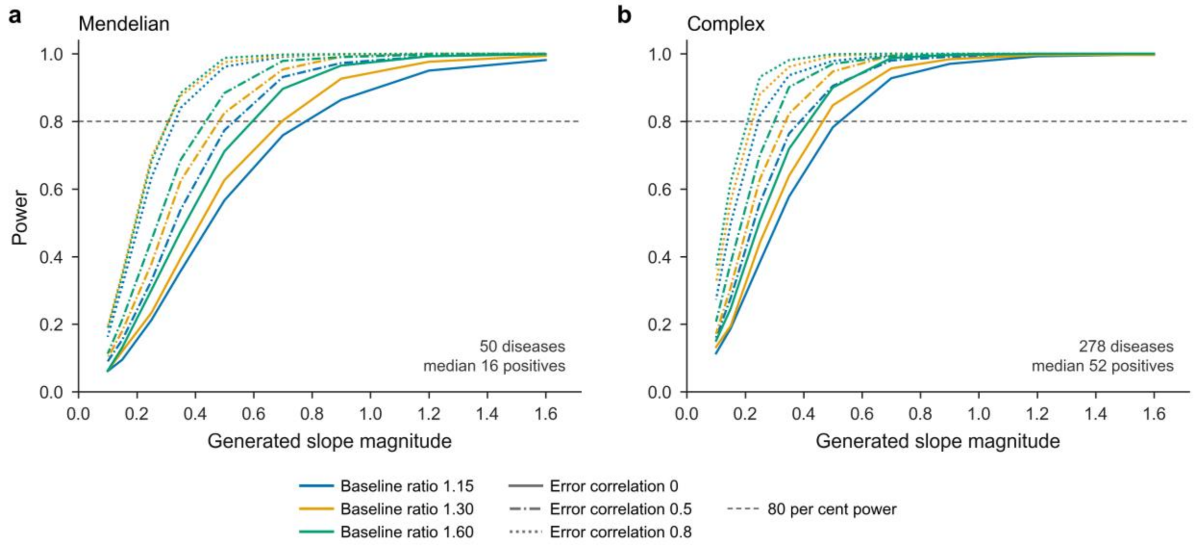


Power against the magnitude of the generated slope of log $R$ on onset, for each baseline ratio and each assumed correlation between the two nested discrimination estimates, on the Mendelian arm (**a**) and the complex arm (**b**). Each panel states the diseases clearing the floor and their median positive count.

**Figure S3** Left truncation in the Finnish registry against the English series

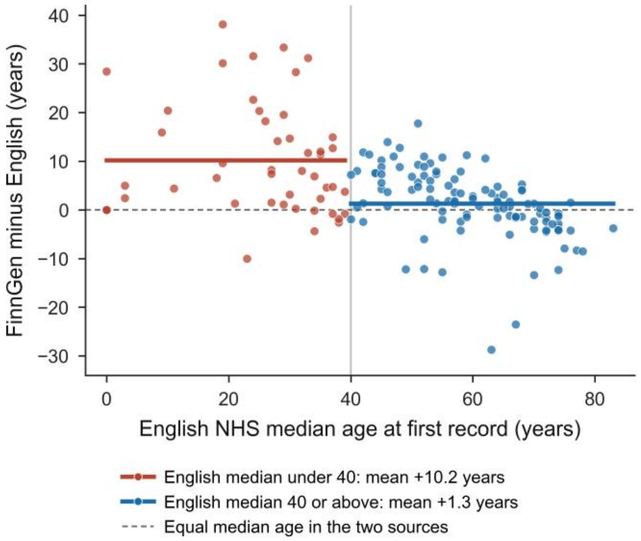


Difference between the Finnish and English median age at first record, against the English median, for 161 conditions matched by name. The horizontal lines are the group means either side of an English median of 40 years.

**Figure S4** Positive rows lost to the autosomal restriction, by onset level

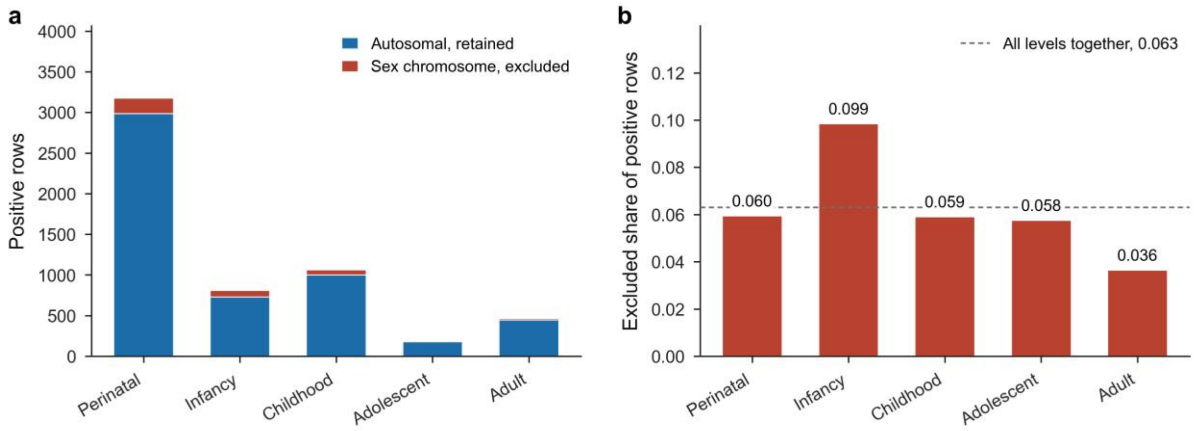


**a** Positive rows retained and excluded at each onset level, counted over the positives whose gene carries both primary scores. **b** The excluded share at each level against the share across all levels.

**Figure S5** Median standardised B2 by onset level

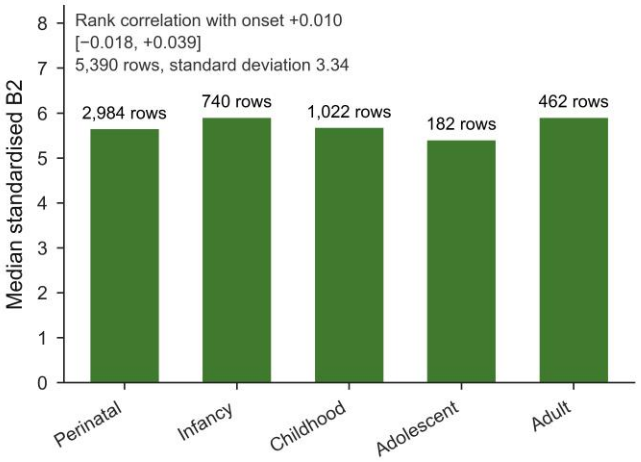


Median standardised B2 for the gene-disease rows at each onset level, with the row count above each bar and the rank correlation with onset given in the panel.

**Figure S6** Evo 2 against DepMap essentiality, with two composition baselines

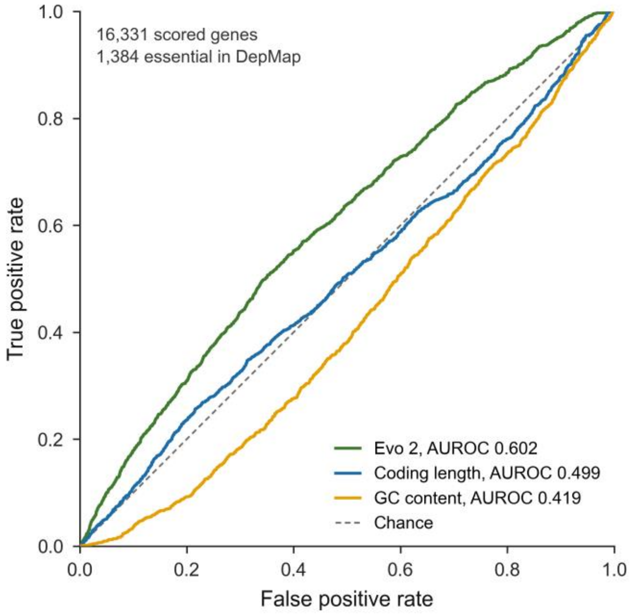


Receiver operating characteristic curves for the Evo 2 stop-codon score and for coding length and GC content on the same genes, with the area under each curve. Evo 2 is ranked as the negative of its score, so a more surprising insertion ranks a gene as more essential.

**Figure S7** Composition of the gene set the insertion rule can score

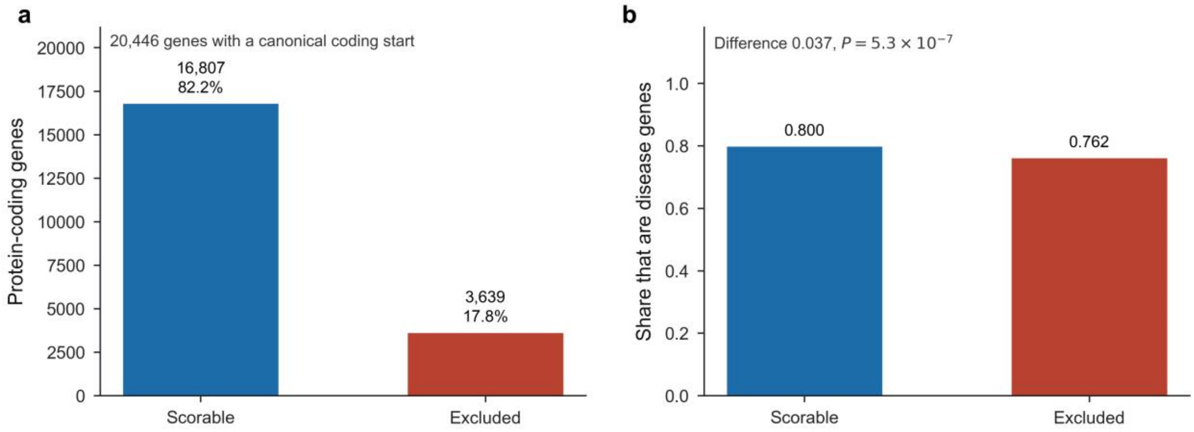


**a** Protein-coding genes with a canonical coding start, split by whether the first coding exon is long enough to hold the furthest insertion. **b** The share of each set that are disease genes, with the difference and its chi-squared *P* value.

**Figure S8** Row sets in use and the unconditioned contrast on each

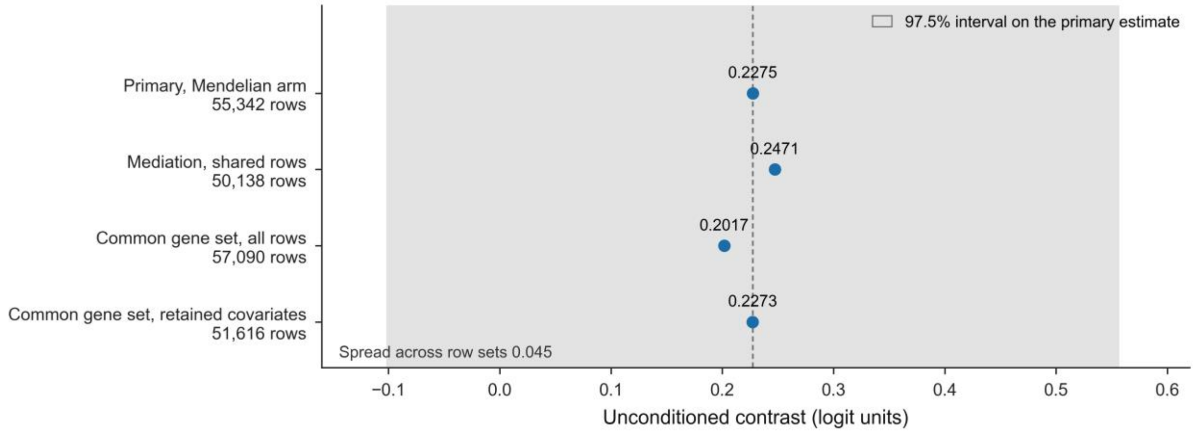


The contrast fitted without any gene-class covariate on each row set, against the 97.5% interval on the primary estimate. The spread across row sets is an order of magnitude below that interval.

#### Additional tables

**Table S1** Data sources, with the data taken from each, the release used, the date of retrieval and the licence under which the source is published.

| Source | Data | Release | Retrieved | Licence |
|---|---|---|---|---|
| Orphanet | Age of onset and inheritance, gene associations, disease cross-references | 1.3.42, 23 June 2026 | 25 August 2026 | CC BY 4.0 |
| Open Targets Platform | Disease index, targets, evidence from Orphanet, GWAS credible sets, ClinVar and gene burden, molecular interactions, GTEx expression, DepMap essentiality | 26.06 | 25 to 26 August 2026 | CC0 1.0 |
| gnomAD | Gene constraint metrics | v4.1 and v2.1.1 | 25 August 2026 | ODbL 1.0 |
| GeneBayes | Per-gene s_het estimates | Zenodo deposit, 19 December 2023 | 25 August 2026 | CC BY 4.0 |
| UCSC Genome Browser | phyloP 241-way, phyloP 100-way and phastCons 100-way tracks | hg38 | 26 August 2026 | UCSC terms of use |
| GENCODE | Gene annotation, protein-coding transcripts, primary assembly | Release 44 | 26 August 2026 | Free with attribution |
| Gene Ontology | Biological-process ontology | 26 July 2026 | 26 August 2026 | CC BY 4.0 |
| Mondo Disease Ontology | Disease ontology | 4 August 2026 | 26 August 2026 | CC BY 4.0 |
| FinnGen, through Risteys | Median age at first recorded event per endpoint | R13 | 26 August 2026 | Risteys terms of use |
| Kuan et al. 2019 | Median age at first record for 308 conditions | Published appendix | 26 August 2026 | CC BY 4.0 |
| Spence et al. 2025 | Per-gene effect estimates across UK Biobank traits | Repository commit d856550 | 26 August 2026 | MIT |
| BetaScan2 | Standardised B2 for the CEU, CHB and YRI populations | Zenodo deposit v1, 3 February 2020 | 26 August 2026 | CC BY 4.0 |

**Table S2** Onset coverage in the Orphanet natural history product.

| Quantity | ORPHAcodes |
|---|---|
| ORPHAcodes in the natural history product | 7,790 |
| With any age-of-onset annotation | 6,982 |
| Annotated only as spanning all ages | 868 |
| With an ordinal onset interval | 6,114 |
| With at least one gene association | 4,245 |
| With both an ordinal interval and a gene | 3,524 |

**Table S3** Orphanet onset intervals before collapse, for the diseases carrying both an ordinal interval and a gene association, with the level each interval is merged into.

| Orphanet interval | Diseases | Collapsed level |
|---|---|---|
| antenatal | 521 | perinatal |
| neonatal | 1,361 | perinatal |
| infancy | 502 | infancy |
| childhood | 663 | childhood |
| adolescent | 120 | adolescent |

| Orphanet interval | Diseases | Collapsed level |
|---|---|---|
| adult | 353 | adult |
| elderly | 4 | adult |
| **total** | **3,524** | |

**Table S4** Composition of the adult-onset Mendelian cell by disease category. Categories are assigned by a keyword pass over the Orphanet preferred name.

| Category | Diseases | Gene-disease pairs |
|---|---|---|
| other | 128 | 240 |
| neurodegenerative | 51 | 96 |
| hereditary cancer predisposition | 49 | 94 |
| metabolic | 29 | 30 |
| cardiac or neuromuscular | 26 | 26 |

**Table S5** Inheritance composition by onset level, counted in diseases. The dominant share is taken over the two resolved modes.

| Onset | Dominant | Recessive | Mixed | Other or unknown | Dominant share |
|---|---|---|---|---|---|
| perinatal | 488 | 1,043 | 138 | 80 | 0.319 |
| infancy | 111 | 315 | 25 | 26 | 0.261 |
| childhood | 213 | 307 | 46 | 35 | 0.410 |
| adolescent | 50 | 38 | 6 | 9 | 0.568 |
| adult | 155 | 62 | 15 | 51 | 0.714 |

**Table S6** Agreement between the two national registries on median age at first recorded event, over conditions matched by name.

| Quantity | Value |
|---|---|
| Conditions matched | 161 |
| Spearman correlation | 0.867 |
| Pearson correlation | 0.870 |
| Median difference, FinnGen minus English | +2.3 years |
| Mean difference where the English median is under 40 | +10.2 years |
| Mean difference where the English median is 40 or above | +1.3 years |
| Conditions under 40 | 48 |
| Conditions at 40 or above | 113 |

**Table S7** Design-stage power simulation, with standard errors clustered on disease alone. The contrast is simulated at 1,000 replicates per cell, the incremental-value slope at 800. The minimum detectable effects reported in the paper are computed from the realised standard errors instead.

| Simulation | Stratum or cell | Minimum detectable effect |
|---|---|---|
| Contrast | All Mendelian, five onset levels, matched controls, score correlation 0.2 | 0.048 AUROC |

| Simulation | Stratum or cell | Minimum detectable effect |
|---|---|---|
| Contrast | All Mendelian, five onset levels, matched controls, score correlation 0.4 | 0.043 AUROC |
| Contrast | All Mendelian, five onset levels, matched controls, score correlation 0.6 | 0.036 AUROC |
| Contrast | All Mendelian, five onset levels, matched controls, score correlation 0.613, measured | 0.035 AUROC |
| Contrast | All Mendelian, five onset levels, random controls, score correlation 0.2 | 0.048 AUROC |
| Contrast | All Mendelian, five onset levels, random controls, score correlation 0.4 | 0.043 AUROC |
| Contrast | All Mendelian, five onset levels, random controls, score correlation 0.6 | 0.035 AUROC |
| Contrast | All Mendelian, five onset levels, random controls, score correlation 0.613, measured | 0.034 AUROC |
| Contrast | All Mendelian, two onset levels, matched controls, score correlation 0.2 | 0.059 AUROC |
| Contrast | All Mendelian, two onset levels, matched controls, score correlation 0.4 | 0.048 AUROC |
| Contrast | All Mendelian, two onset levels, matched controls, score correlation 0.6 | 0.042 AUROC |
| Contrast | All Mendelian, two onset levels, matched controls, score correlation 0.613, measured | 0.041 AUROC |
| Contrast | All Mendelian, two onset levels, random controls, score correlation 0.2 | 0.057 AUROC |
| Contrast | All Mendelian, two onset levels, random controls, score correlation 0.4 | 0.048 AUROC |
| Contrast | All Mendelian, two onset levels, random controls, score correlation 0.6 | 0.042 AUROC |
| Contrast | All Mendelian, two onset levels, random controls, score correlation 0.613, measured | 0.041 AUROC |
| Contrast | Dominant only, five onset levels, matched controls, score correlation 0.2 | 0.078 AUROC |
| Contrast | Dominant only, five onset levels, matched controls, score correlation 0.4 | 0.070 AUROC |
| Contrast | Dominant only, five onset levels, matched controls, score correlation 0.6 | 0.058 AUROC |
| Contrast | Dominant only, five onset levels, matched controls, score correlation 0.613, measured | 0.052 AUROC |
| Contrast | Dominant only, five onset levels, random controls, score correlation 0.2 | 0.076 AUROC |
| Contrast | Dominant only, five onset levels, random controls, score correlation 0.4 | 0.069 AUROC |
| Contrast | Dominant only, five onset levels, random controls, score correlation 0.6 | 0.055 AUROC |
| Contrast | Dominant only, five onset levels, random controls, score correlation 0.613, measured | 0.050 AUROC |
| Contrast | Dominant only, two onset levels, matched controls, score correlation 0.2 | 0.078 AUROC |
| Contrast | Dominant only, two onset levels, matched controls, score correlation 0.4 | 0.071 AUROC |
| Contrast | Dominant only, two onset levels, matched controls, score correlation 0.6 | 0.060 AUROC |
| Contrast | Dominant only, two onset levels, matched controls, score correlation 0.613, measured | 0.054 AUROC |

| Simulation | Stratum or cell | Minimum detectable effect |
|---|---|---|
| Contrast | Dominant only, two onset levels, random controls, score correlation 0.2 | 0.077 AUROC |
| Contrast | Dominant only, two onset levels, random controls, score correlation 0.4 | 0.070 AUROC |
| Contrast | Dominant only, two onset levels, random controls, score correlation 0.6 | 0.057 AUROC |
| Contrast | Dominant only, two onset levels, random controls, score correlation 0.613, measured | 0.051 AUROC |
| Incremental value, Mendelian arm | true ratio 1.15, correlation between nested AUROCs 0.0 | 0.779 slope |
| Incremental value, Mendelian arm | true ratio 1.15, correlation between nested AUROCs 0.5 | 0.533 slope |
| Incremental value, Mendelian arm | true ratio 1.15, correlation between nested AUROCs 0.8 | 0.331 slope |
| Incremental value, Mendelian arm | true ratio 1.3, correlation between nested AUROCs 0.0 | 0.697 slope |
| Incremental value, Mendelian arm | true ratio 1.3, correlation between nested AUROCs 0.5 | 0.481 slope |
| Incremental value, Mendelian arm | true ratio 1.3, correlation between nested AUROCs 0.8 | 0.309 slope |
| Incremental value, Mendelian arm | true ratio 1.6, correlation between nested AUROCs 0.0 | 0.596 slope |
| Incremental value, Mendelian arm | true ratio 1.6, correlation between nested AUROCs 0.5 | 0.436 slope |
| Incremental value, Mendelian arm | true ratio 1.6, correlation between nested AUROCs 0.8 | 0.309 slope |
| Incremental value, complex arm | true ratio 1.15, correlation between nested AUROCs 0.0 | 0.524 slope |
| Incremental value, complex arm | true ratio 1.15, correlation between nested AUROCs 0.5 | 0.389 slope |
| Incremental value, complex arm | true ratio 1.15, correlation between nested AUROCs 0.8 | 0.244 slope |
| Incremental value, complex arm | true ratio 1.3, correlation between nested AUROCs 0.0 | 0.466 slope |
| Incremental value, complex arm | true ratio 1.3, correlation between nested AUROCs 0.5 | 0.338 slope |
| Incremental value, complex arm | true ratio 1.3, correlation between nested AUROCs 0.8 | 0.225 slope |
| Incremental value, complex arm | true ratio 1.6, correlation between nested AUROCs 0.0 | 0.417 slope |
| Incremental value, complex arm | true ratio 1.6, correlation between nested AUROCs 0.5 | 0.300 slope |
| Incremental value, complex arm | true ratio 1.6, correlation between nested AUROCs 0.8 | 0.208 slope |

**Table S8** Row sets in use, with what each holds and the unconditioned contrast fitted on it.

| Row set | What it holds | Rows | Contrast |
|---|---|---|---|
| Primary, Mendelian arm | Every gene carrying s_het and the conservation metric | 55,342 | 0.2275 |
| Mediation, shared rows | The rows every mediation specification can be fitted on | 50,138 | 0.2471 |

| Row set | What it holds | Rows | Contrast |
|---|---|---|---|
| Common gene set, all rows | Only genes carrying all three constraint metrics | 57,090 | 0.2017 |
| Common gene set, retained covariates | As above, restricted to the three retained covariates | 51,616 | 0.2273 |
| Common gene set, all four covariates | As above, including disease breadth | 51,585 | 0.2295 |

**Table S9** Change in the contrast under conditioning, by covariate and constraint metric, over 400 resamples of diseases on a fixed row set.

| Covariate | s_het | LOEUF | Missense Z |
|---|---|---|---|
| Haploinsufficiency (pLI) | +0.000 [-0.010, +0.022] | +0.035 [+0.021, +0.055] | +0.024 [+0.013, +0.045] |
| Developmental share | +0.007 [-0.007, +0.038] | -0.013 [-0.020, +0.004] | +0.002 [-0.024, +0.015] |
| Trait breadth | -0.001 [-0.005, +0.005] | -0.002 [-0.004, +0.001] | -0.002 [-0.005, +0.005] |
| Disease breadth | +0.054 [+0.019, +0.087] | +0.032 [-0.004, +0.044] | +0.019 [-0.013, +0.046] |
| The three retained, jointly | +0.006 [-0.006, +0.049] | +0.020 [+0.009, +0.050] | +0.025 [-0.002, +0.052] |
| All four together | +0.046 [+0.020, +0.101] | +0.048 [+0.011, +0.078] | +0.034 [-0.001, +0.081] |

**Table S10** Rank correlation between each adjustment covariate and onset, among disease genes and across all design rows, resampled over diseases.

| Covariate | Stratum | Rows | Spearman | 95% interval |
|---|---|---|---|---|
| Developmental share | disease genes | 5,251 | -0.0825 | -0.1211 to -0.0477 |
| Developmental share | all design rows | 50,451 | -0.0204 | -0.0313 to -0.0092 |
| Haploinsufficiency (pLI) | disease genes | 5,321 | +0.0145 | -0.0340 to +0.0623 |
| Haploinsufficiency (pLI) | all design rows | 55,340 | +0.0060 | -0.0060 to +0.0170 |
| Trait breadth | disease genes | 5,262 | +0.0111 | -0.0153 to +0.0373 |
| Trait breadth | all design rows | 54,980 | +0.0033 | -0.0062 to +0.0124 |
| Disease breadth | disease genes | 5,322 | -0.0106 | -0.0412 to +0.0232 |
| Disease breadth | all design rows | 55,176 | +0.0064 | -0.0112 to +0.0278 |

**Table S11** The contrast under the two-way specification and under the three-way specification that lets the onset slope vary by mode of inheritance. Both columns are fitted on the genes carrying all three constraint metrics, so the two specifications are compared on one sample.

| Constraint metric | Two-way | 97.5% interval | Three-way | 97.5% interval |
|---|---|---|---|---|
| s_het | 0.202 | -0.136 to 0.539 | 0.465 | -0.207 to 1.136 |
| Missense Z | 0.375 | 0.134 to 0.617 | 0.479 | 0.056 to 0.903 |
| LOEUF | 0.481 | 0.297 to 0.664 | 0.401 | -0.093 to 0.895 |

**Table S12** Rank correlation among the constraint metrics within each onset level. LOEUF is inverted by construction, so a negative value against it is agreement.

| Stratum | Genes | s_het and LOEUF | s_het and missense Z | LOEUF and missense Z |
|---|---|---|---|---|
| all genes | 3,788 | -0.821 | 0.626 | -0.733 |
| perinatal | 2,079 | -0.807 | 0.596 | -0.722 |

| Stratum | Genes | s_het and LOEUF | s_het and missense Z | LOEUF and missense Z |
|---|---|---|---|---|
| infancy | 678 | -0.878 | 0.717 | -0.749 |
| childhood | 848 | -0.820 | 0.656 | -0.738 |
| adolescent | 172 | -0.755 | 0.600 | -0.753 |
| adult | 356 | -0.816 | 0.645 | -0.743 |

**Table S13** Discrimination of Mendelian disease genes within each inheritance stratum, against the pool of scored genes and against matched controls, over 1,000 resamples of genes.

| Stratum | Metric | Disease genes | Pool | AUROC | 95% interval |
|---|---|---|---|---|---|
| recessive | s_het | 2,173 | 17,434 | 0.4872 | 0.4763 to 0.4979 |
| recessive | LOEUF | 2,173 | 17,434 | 0.5334 | 0.5215 to 0.5438 |
| recessive | Missense Z | 2,173 | 17,434 | 0.5212 | 0.5089 to 0.5342 |
| dominant | s_het | 1,218 | 17,434 | 0.7229 | 0.7071 to 0.7372 |
| dominant | LOEUF | 1,218 | 17,434 | 0.7254 | 0.7095 to 0.7404 |
| dominant | Missense Z | 1,218 | 17,434 | 0.6815 | 0.6652 to 0.6975 |

Paired differences on the same resamples, against the pool.

| Stratum | Comparison | Difference | 95% interval | Excludes zero |
|---|---|---|---|---|
| recessive | LOEUF minus s_het | +0.0462 | 0.0374 to 0.0554 | yes |
| recessive | Missense Z minus s_het | +0.0341 | 0.0230 to 0.0462 | yes |
| dominant | LOEUF minus s_het | +0.0025 | -0.0075 to 0.0123 | no |
| dominant | Missense Z minus s_het | -0.0414 | -0.0576 to -0.0272 | yes |

Against ten controls per gene, matched on coding sequence length decile, expected loss-of-function count decile and GC decile.

| Stratum | Metric | Disease genes | Controls | AUROC | 95% interval |
|---|---|---|---|---|---|
| recessive | s_het | 2,173 | 20,908 | 0.4572 | 0.4471 to 0.4688 |
| recessive | LOEUF | 2,173 | 20,908 | 0.4505 | 0.4405 to 0.4602 |
| recessive | Missense Z | 2,173 | 20,908 | 0.4722 | 0.4616 to 0.4833 |
| dominant | s_het | 1,218 | 11,751 | 0.6742 | 0.6575 to 0.6896 |
| dominant | LOEUF | 1,218 | 11,751 | 0.6499 | 0.6335 to 0.6643 |
| dominant | Missense Z | 1,218 | 11,751 | 0.6336 | 0.6182 to 0.6475 |

Paired differences against matched controls, on the same resamples.

| Stratum | Comparison | Difference | 95% interval | Excludes zero |
|---|---|---|---|---|
| recessive | LOEUF minus s_het | -0.0067 | -0.0136 to -0.0004 | yes |
| recessive | Missense Z minus s_het | +0.0150 | 0.0049 to 0.0246 | yes |
| dominant | LOEUF minus s_het | -0.0243 | -0.0323 to -0.0161 | yes |
| dominant | Missense Z minus s_het | -0.0406 | -0.0549 to -0.0265 | yes |

Covariate balance, as medians and as the standardised mean difference of the disease genes against each reference set.

| Stratum | Covariate | Disease genes | Matched controls | Pool | SMD, matched | SMD, pool |
|---|---|---|---|---|---|---|
| recessive | Coding sequence length | 1,662 | 1,665 | 1,359 | 0.074 | 0.309 |
| recessive | Expected loss-of-function count | 50.2 | 50.4 | 35.4 | 0.097 | 0.404 |
| recessive | GC content | 0.528 | 0.528 | 0.530 | 0.006 | -0.042 |
| dominant | Coding sequence length | 1,851 | 1,833 | 1,359 | 0.098 | 0.510 |
| dominant | Expected loss-of-function count | 54.7 | 56.2 | 35.4 | 0.103 | 0.566 |
| dominant | GC content | 0.544 | 0.547 | 0.530 | 0.006 | 0.132 |

**Table S14** Incremental value of the evolutionary scores by onset bin, on both arms. R is the ratio of the lift over chance of a model with the evolutionary scores to one without them.

| Arm | Onset | Diseases | Median R | First quartile | Third quartile |
|---|---|---|---|---|---|
| Mendelian | 0.000 | 31 | 1.161 | 0.999 | 1.282 |
| Mendelian | 0.250 | 4 | 1.283 | 1.155 | 1.353 |
| Mendelian | 0.500 | 9 | 1.292 | 0.956 | 1.525 |
| Mendelian | 0.750 | 3 | 1.023 | 0.990 | 1.415 |
| Mendelian | 1.000 | 3 | 1.055 | 0.899 | 1.144 |
| Complex | 0.000 | 2 | 1.723 | 1.667 | 1.780 |
| Complex | 0.143 | 7 | 1.320 | 1.137 | 1.467 |
| Complex | 0.286 | 11 | 1.161 | 1.003 | 1.325 |
| Complex | 0.429 | 33 | 1.241 | 1.071 | 1.360 |
| Complex | 0.571 | 36 | 1.213 | 1.108 | 1.304 |
| Complex | 0.714 | 90 | 1.259 | 1.116 | 1.387 |
| Complex | 0.857 | 76 | 1.199 | 1.035 | 1.420 |
| Complex | 1.000 | 23 | 1.258 | 1.103 | 1.508 |

**Table S15** Evo 2 scoring against the published human essentiality benchmark, and the composition of the gene set the insertion rule can score. The Evo 2, coding length and GC content rows are rank statistics of a single score. The two combined rows are out-of-fold discrimination of a fitted logistic model over five folds, so they are not an upper bound on the rows above: on the same rows the out-of-fold figure for Evo 2 alone is 0.601, and for GC content alone, which ranks below chance as a raw score, it is 0.580.

| Quantity | Value |
|---|---|
| Genes scored and carrying a DepMap label | 16,331 |
| Essential genes among them | 1,384 |
| Evo 2 AUROC | 0.6018 |
| Coding length alone | 0.4991 |
| GC content alone | 0.4193 |
| Both baselines together | 0.5800 |
| Baselines plus Evo 2 | 0.5830 |
| Published reference | 0.66 |

| Quantity | Value |
|---|---|
| Difference from the reference | 0.0582 |
| Inclusion criterion | within 0.03 of the reference |
| Criterion met | no |

Gene set the insertion rule excludes.

| Quantity | Value |
|---|---|
| Protein-coding genes with a canonical coding start | 20,446 |
| Scorable under the insertion rule | 16,807 |
| Excluded for a short first coding exon | 3,639 |
| Excluded share | 0.1780 |
| Share carrying disease evidence in any arm, scorable | 0.7996 |
| Share carrying disease evidence in any arm, excluded | 0.7623 |
| Difference | 0.0373 |

**Table S16** Each score family's main effect and onset decay, with 97.5% intervals from the two-way cluster-robust covariance. The decay is the negative of the score-by-onset coefficient, so a positive value is a loss of discriminative power as onset rises. Γ is the difference between the two decays. The third pair drops the Orphanet associations recording somatic mutations and those not yet assessed.

| Specification | Family | Main effect | 97.5% interval | Onset decay | 97.5% interval | Decay / main |
|---|---|---|---|---|---|---|
| Primary, unconditioned | Conservation | 0.263 | 0.120 to 0.406 | 0.360 | 0.180 to 0.539 | 1.37 |
| Primary, unconditioned | Constraint | 0.212 | 0.005 to 0.420 | 0.132 | −0.061 to 0.326 | 0.62 |
| Three retained covariates | Conservation | 0.183 | 0.025 to 0.341 | 0.396 | 0.210 to 0.583 | 2.16 |
| Three retained covariates | Constraint | 0.233 | 0.013 to 0.452 | 0.123 | −0.074 to 0.320 | 0.53 |
| Somatic and unassessed dropped | Conservation | 0.292 | 0.146 to 0.437 | 0.285 | 0.098 to 0.471 | 0.98 |
| Somatic and unassessed dropped | Constraint | 0.099 | −0.127 to 0.324 | 0.252 | 0.050 to 0.454 | 2.56 |
